# Rett syndrome lifespan extension in mice via AI-guided ADAR editing

**DOI:** 10.64898/2026.06.23.734060

**Authors:** Yiannis A. Savva, Brian J. Booth, Lucia Shumaker, Rachael Fasnacht, Stephen M. Burleigh, Yue Jiang, Yingxin Cao, Brian Johnson, Lina R. Bagepalli, Forrest Golic, Nicole Enger, Rachel Feiring, Andrew Sadowski, Scott Rich, Anupama Lakshmanan, Nikita Milani, Eric M. Chadwick, Collin Hauskins, Melissa G. Works, David J. Huss, Adrian W. Briggs, Alison A. VanSchoiack

**Affiliations:** Shape Therapeutics, 700 Dexter Avenue N, 98109 Seattle WA

## Abstract

Rett syndrome is a severe neurodevelopmental disorder primarily caused by mutations in the *MECP2* gene. A significant subset of severe cases are driven by nonsense mutations that generate premature stop codons, leading to loss of functional MeCP2 protein. Here, we describe a novel therapeutic strategy that uses endogenous adenosine deaminase acting on RNA (ADAR) enzymes to correct the R168X mutation at the RNA level. Using generative artificial intelligence trained on large empirical datasets, we engineered guide RNAs (gRNAs) that recruit endogenous ADAR to convert the mutant stop codon (UGA) into a tryptophan (UGG) to restore full-length MeCP2. Once incorporated into an optimized expression system based on endogenous small nuclear RNA regulatory elements and packaged into adeno-associated virus, these gRNAs enabled precise RNA editing at the target site with minimal off-target activity across the transcriptome while restoring full-length MeCP2 protein expression in patient-derived induced pluripotent stem cell neurons. Delivered intravenously to an R168X mouse model, the gRNAs achieved ~70% targeted RNA editing and substantially restored MeCP2 throughout the brain resulting in markedly improved Rett-like phenotypes and significantly extended lifespan. These findings demonstrate that AI-guided ADAR-mediated RNA editing is a precise and efficient technology for correcting nonsense mutations, with therapeutic potential for Rett syndrome and other genetic diseases.

---

Rett syndrome (RTT) is a severe neurodevelopmental disorder predominantly resulting from loss-of-function mutations in the X-linked *MECP2* gene, which encodes the methyl-CpG-binding protein 2, a critical transcriptional regulator^1^. The R168X nonsense mutation, a common and highly deleterious variant, introduces a premature UGA stop codon that prevents formation of functional MeCP2 protein^2^. This disruption impairs transcriptional regulation essential for neuronal development and function, leading to severe neurological manifestations, including developmental regression, motor dysfunction, and cognitive deficits^3^. Due to X-linked inheritance, males exhibit more severe phenotypes, often resulting in premature mortality^4^. Current therapeutic approaches for RTT are predominantly symptomatic and lack interventions that address the underlying genetic etiology^5^. By leveraging ADAR mediated RNA editing to correct the *MECP2* transcript, protein function can be restored within the bounds of endogenous regulation, preventing the risks of transgene overexpression^6,7^. In the context of RTT’s X-linked mosaicism, the ADAR correction approach provides a particularly elegant solution since diseased cells undergo correction while wild-type (WT) cells remain stable under a synonymous change, ensuring the delicate protein balance necessary for cellular function^8,9^.

Multiple emerging therapeutic strategies have the potential to target the R168X premature stop codon mutation at distinct molecular levels. At the DNA level, CRISPR-Cas9 genome editing can directly and permanently correct the pathogenic variant^10^, though it carries risks of off-target genomic alterations and faces considerable hurdles in delivering large editing complexes safely and efficiently to post-mitotic neurons in the Central Nervous System (CNS)^11^. Adenine base editors (ABEs) offer a more refined alternative, coupling a Cas9 nickase to an adenosine deaminase to achieve precise single-nucleotide correction without generating double-strand breaks, which reduces the risk of insertions or deletions^12^. However, ABEs remain susceptible to off-target DNA and RNA editing and share the delivery limitations of large ribonucleoprotein complexes^13–15^. At the level of protein translation, nonsense suppression strategies, including small-molecule read-through agents such as ataluren (PTC124) and aminoglycoside derivatives like gentamicin, promote ribosomal decoding of the UGA stop codon to generate full-length MeCP2^16^. Their utility may be limited, however, by low read-through efficiency, the potential incorporation of near-cognate amino acids that may impair protein function, and risks of off-target translational effects and systemic toxicity with prolonged use^17^. Finally, adeno-associated virus (AAV) mediated gene replacement therapy can deliver an exogenous functional *MECP2* transgene to restore protein expression. While the *MECP2* gene fits within AAV’s capacity, the requirement for complex regulatory circuits to prevent dosage-related toxicity complicates vector architecture^8^.

RNA editing platforms using chemically modified or genetically encoded ADAR-recruiting guide RNAs (gRNAs) can modulate MECP2 transcript processing or directly recode the premature stop codon at the RNA level^18,19^. These strategies offer a potentially safer route than DNA editing by avoiding permanent genomic changes, and their use of an endogenous editing enzyme drastically reduces immunogenicity risks from foreign proteins. Custom editing with endogenous ADAR has also been shown to cause only minimal transcriptome-wide off-target effects compared to DNA editors^20^. Unlike DNA editing systems, RNA editing with ADAR works particularly well in post-mitotic cells like neurons, which are critical target cells in RTT. While directly delivered chemically modified oligonucleotides that recruit ADAR have shown promise in the clinic in liver-targeted therapies, they require regular redosing and their delivery to the brain remains a major challenge. In contrast, AAV presents a highly attractive option to deliver gRNAs to the brain, given AAV’s proven ability to deeply penetrate brain tissue^21^, the potential long-term safety advantage of gRNAs being comprised of fully natural RNA, and the promise of a one-time, single-dose regimen for serious diseases like RTT.

In this study, we utilized generative artificial intelligence (AI) models trained on large empirical datasets to computationally design and optimize ADAR-recruiting gRNAs exhibiting high editing efficiency and target specificity^22^, while also identifying gRNAs that are allele specific or allele cross-reactive and species cross-reactive, simplifying the pre-clinical path. The leading gRNA candidates were incorporated into a published small nuclear RNA (snRNA) expression cassette^23^ that we further optimized with improved 5′ and 3′ regulatory elements alongside a novel architectural configuration. Performance was rigorously evaluated for efficacy and specificity in correcting the R168X nonsense mutation using both *in vitro* and *in vivo* disease-relevant models. Notably, AAV-mediated delivery of these snRNA-gRNA pay-loads to the CNS of adult RTT mouse model resulted in unprecedented levels of precise, endogenous ADAR-mediated A-to-I RNA editing. This editing substantially restored functional MeCP2 protein expression in the mature brain, concomitant with amelioration of RTT-associated behavioral and physiological phenotypes, as well as a marked extension of lifespan. Critically, no detectable off-target editing was observed at the transcriptomic level *in vitro* and the snRNA-gRNA payloads were well-tolerated *in vivo*. These results advance RNA editing-based therapeutic modalities for RTT and establish a versatile framework for addressing other genetic disorders amenable to targeted, endogenous ADAR-driven RNA correction.

## Precision AI engineering of robust ADAR-recruiting gRNAs

ADAR-mediated RNA editing of the R168X nonsense mutation recodes the premature UGA stop codon to UGG, resulting in tryptophan incorporation rather than restoration of the wild-type (WT) arginine residue. To determine whether this amino acid substitution produces a stable, full-length MeCP2 protein capable of recapitulating WT subcellular localization, we rationally designed an ADAR-recruiting gRNA (MECP2-01; **Supplementary Fig. 1a**) to test across multiple *in vitro* systems. We first assessed editing efficiency in both human and mouse HEK293 minigene systems. Although this design achieved substantial editing at the target adenosine on both the mouse and human transcripts, it lacked positional specificity, with bystander editing detected at the +5 and +8 positions relative to the target site **(Supplementary Fig. 1b)**.

To enable delivery to neurons, the gRNA was incorporated into a previously described snRNA expression cassette^23^ and packaged into AAV-PHP.eB capsid for evaluation in RTT patient derived induced pluripotent stem cell (iPSC) neurons and primary neurons cultured from Mecp2 R168X male mice. Transduction at both low (5 × 10^4^ vg/cell) and high (5 × 10^5^ vg/cell) doses produced dose-dependent RNA editing at the target adenosine **(Supplementary Fig. 1c)**. Immunocytochemistry confirmed restoration of MeCP2 protein expression in both cell types (**Supplementary Fig. 1d–e**). Critically, the restored protein localized to the nucleus, similar to wild-type MeCP2 and consistent with MeCP2’s established function as an epigenetic regulator and chromatin-associated protein. Collectively, these findings demonstrate that A-to-I editing-mediated recoding of the pathogenic UGA stop codon to UGG constitutes a viable therapeutic strategy for this class of Rett syndrome variants. We therefore hypothesized that an optimized gRNA, with improved editing efficiency and target specificity beyond that achieved by rational design, could direct precise adenosine deamination within the UGA codon, enabling robust restoration of functional MeCP2 protein and downstream functional phenotypes.

To achieve this, we employed DeepREAD, a bit-diffusion deep learning model trained to generate ADAR-compatible double-stranded RNA substrates optimized for high editing efficiency and specificity^22^ in targeting diverse codons across human and mouse *MECP2* sequences **(Supplementary Fig. 2a)**. The DeepREAD-directed gRNA design proceeds through sequential, discrete stages (**Supplementary Fig. 2b**). This model was trained on data from a biochemical assay featuring shorter dsRNA structures (~41 nucleotides), reflecting practical screening limitations that precluded use of longer substrates; accordingly, it generates *micro-footprints* - ~41-nucleotide antisense gRNA segments incorporating secondary structural features predicted to promote ADAR specificity and target adenosine deamination. To further improve RNA editing efficiency, micro-footprints are integrated into extended gRNA backbones (~100 nucleotides) featuring strategically positioned symmetrical 6/6 nucleotide loops, termed *barbells*, relative to the target site. The resulting full-length sequence is termed the *macro-footprint*. Using DeepREAD, we generated micro-footprint variants in two categories: a) allele-specific gRNAs, designed to preferentially direct editing to the mutant R168X allele while minimizing activity on the WT allele, reflecting the heterozygous state in female RTT patients. b) Crossreactive gRNAs compatible with both human and mouse target sequences, accounting for three nucleotide differences between species that may influence dsRNA structure formation upon gRNA hybridization (**Supplementary Fig. 2c**).

Selected micro-footprints were appended to three distinct macro-footprint backbones and evaluated by transient transfection in **HEK293T-MECP2-4iso** cells harboring cognate minigene reporters (see **Methods**). One macro-footprint configuration (barbell coordinates −15, +33 nucleotide coordinates relative to the target adenosine) consistently supported superior editing efficiency and specificity across tested gRNAs (**Supplementary Fig. 2d**) and was used for all subsequent gRNA designs. Transient transfection and quantitative editing analysis of minigene constructs identified the gRNA MECP2-02 as exhibiting robust allelic discrimination between the mutant and WT transcripts and MECP2-03 and MECP2-04 gRNAs that exhibited allele cross reactivity, while all three exhibited species cross reactivity and a high degree of specificity with minimal bystander editing at adjacent adenosines (**Fig. 1a-b**). Interestingly, despite the structural diversity exhibited by these gRNAs the use of a G at position −2 in MECP2-03 and -04 allows for a canonical or wobble base pairing against R168X and WT transcripts, respectively, enabling the stable formation of the GA-GC bulge at the target adenosine in both alleles, a structure known to facilitate editing of 5’ G targets^24^ (**Fig. 1c)**. While the use of an A at position −2 in MECP2-02 enables the GA-GC bulge only for the R168X transcript, the structure collapses to form an A bulge at the target site in the WT transcript, a structure known to be refractory to editing^25^, thus elucidating a novel molecular switch to discriminate between the two alleles (**Supplementary Fig. 3**). Collectively, these findings indicate that the DeepREAD generative model effectively predicts gRNAs with tunable functional properties and that the outlined engineering workflow enables identification of high-performance gRNAs for further testing in patient-derived iPSC neurons and *in vivo*. Given that editing of the WT allele at this position results in a synonymous codon change (**Supplementary Fig. 2a)**, we reasoned that WT allele editing by this approach in patients is low risk, while enabling pre-clinical studies in healthy non-human primates, and therefore elected to advance all three gRNAs, MECP2-02, MECP2-03, and MECP2-04, forward to derisk any unexpected alterations in efficiency or specificity and to determine if the novel molecular switch for allele specificity or cross reactivity persists in relevant model systems.

**Fig. 1:**
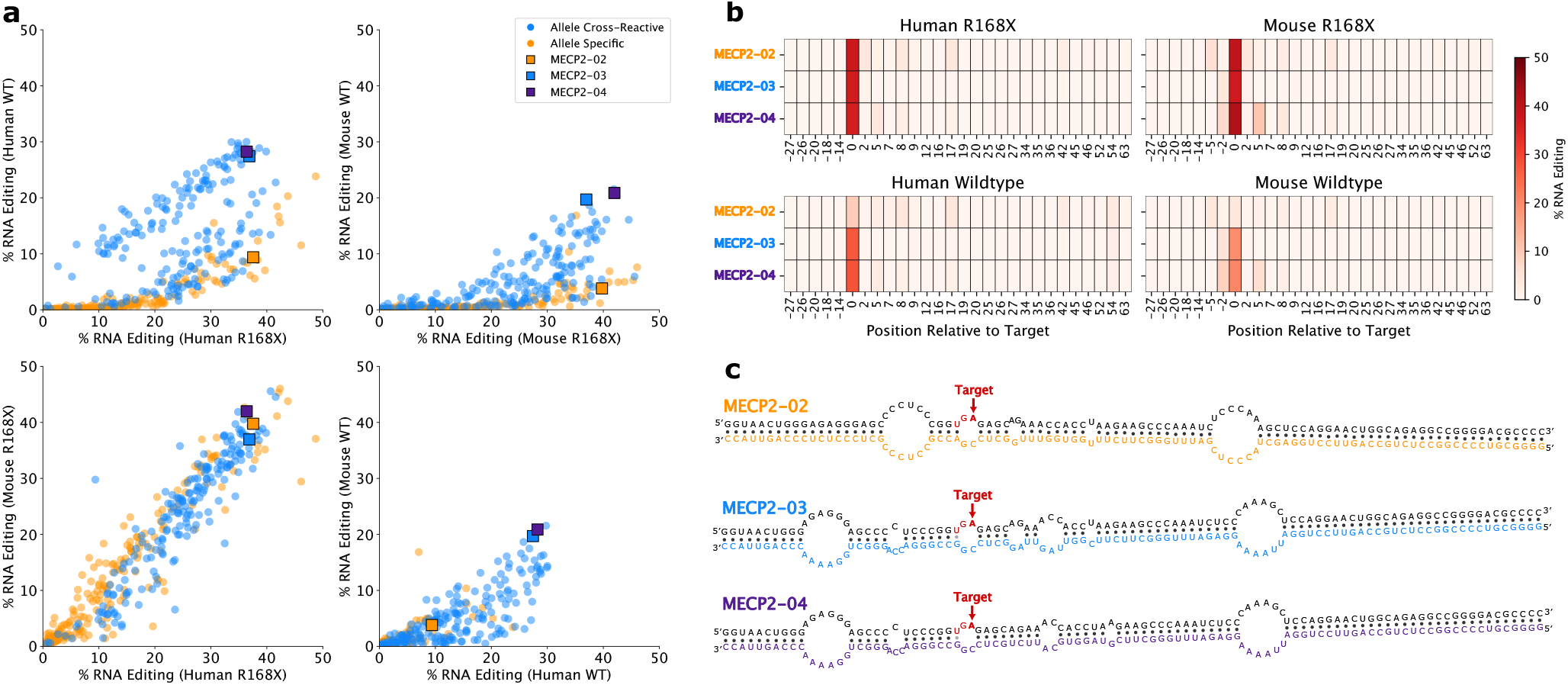
Identification and characterization of lead gRNAs. **a**, On-target A-to-I RNA editing efficiency (%) across four allele/species combinations (Human R168X vs. Human WT; Mouse R168X vs. Mouse WT; cross-species R168X; cross-species WT) for all gRNAs screened in HEK293-MeCP2-4iso cells. Each point represents one gRNA; allele-specific (orange) and allele-cross-reactive (blue) design classes are shown. Lead candidates MECP2-02, MECP2-03, and MECP2-04 are highlighted as squares. **b**, Per-position mean A-to-I editing heatmap across the hybridization region for lead gRNAs (MECP2-02, MECP2-03, and MECP2-04) across four allele/species targets. Rows correspond to gRNA payloads; columns indicate nucleotide position relative to the target adenosine (position 0). Color intensity reflects mean editing percentage (0–50%). **c**, Predicted RNA secondary structures of target–gRNA duplexes for all three lead candidates paired with the Human R168X transcript, computed using Vienna fold RNA duplex. Structures were rendered using VARNA and post-processed to high-light base-pair types (black = canonical; grey = wobble). The R168X codon is shown in red text and the arrow indicates the target adenosine.

### snRNA scaffold optimization for enhanced gRNA expression

Having established a robust computational pipeline for gRNA design and functional prediction, we sought to improve the AAV expression framework to maximize editing efficiency of the MECP2 gRNAs with AAV delivery. Building on prior work demonstrating approximately 75% CNS editing efficiency in a mouse model of Hurler syndrome with an enhanced U7 snRNA cassette^23^, we sought to further improve gRNA expression and editing outcomes through systematic optimization of the 5’ and 3’ cis-regulatory elements flanking the snRNA scaffold **(Fig. 2a)**. Identification of alternative snRNA cassette elements with equivalent activity also provides the sequence heterogeneity necessary to mitigate inter-cassette recombination in multi-copy vector architectures. Using a luciferase reporter assay **(Supplementary Fig. 4a)**, we evaluated candidate core regulatory elements, including the distal sequence element (DSE), proximal sequence element (PSE), the 3’ box and identified configurations yielding approximately two-fold improvements in gRNA expression. These optimized elements were incorporated into a revised expression cassette designated mU7v2.0 **(Supplementary Fig. 5a)**.

**Fig. 2:**
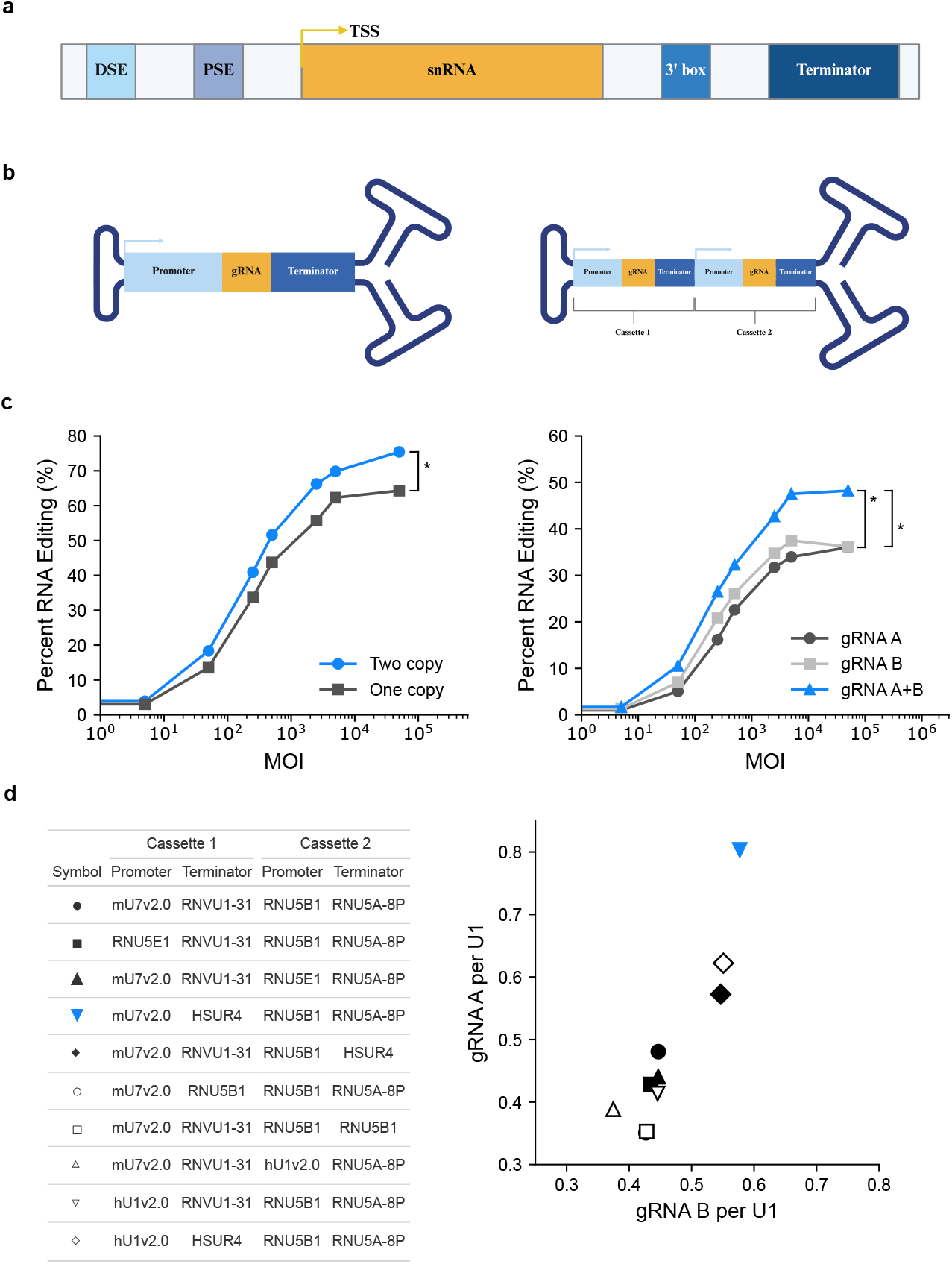
Design and optimization of snRNA promoter-driven gRNA expression cassettes. **a**, Schematic of the snRNA gene regulatory architecture used to drive gRNA expression. The cassette includes a distal sequence element (DSE) and proximal sequence element (PSE) upstream of the transcription start site (TSS), followed by the snRNA coding region (here replaced by the gRNA sequence), a 3’ box, and a transcriptional terminator. Created in BioRender. Vanschoiack, A. (2026) https://BioRender.com/hr59py0. **b**, Schematic representation of AAV vector architectures encoding one (left) or two (right) gRNA expression cassettes. Each cassette consists of an snRNA promoter (blue), gRNA sequence (orange), and terminator sequence (dark blue). The dual-cassette vector contains Cassette 1 and Cassette 2 encoding two copies of a given gRNA molecule. Created in BioRender. Vanschoiack, A. (2026) https://BioRender.com/m159m2c. **c**, Editing efficiency of transduced HEK293 cells, as a function of multiplicity of infection (MOI). (Left) Comparison of one-copy versus two-copy AAV vector configurations across MOIs. Two-sided paired Wilcoxon signed-rank test across 8 MOI levels, p = 0.018. (Right) Editing efficiency of individual gRNAs (gRNA A and gRNA B) compared to the dual-gRNA combination (gRNA A+B). Two-sided paired Wilcoxon signed-rank across 8 MOI levels with Holm correction; A+B vs A and A+B vs B both p = 0.016); *: p < 0.05. **d**, Table listing the promoter (mU7v2.0, RNU5E1, or hU1v2.0) and terminator (RNVU1-31, RNU5B1, RNU5A-8P, or HSUR4) elements used in Cassettes 1 and 2 for each construct, denoted by distinct symbols. Systematic evaluation of top promoter and terminator combinations for dual-cassette gRNA expression, in ARPE-19 cells at 1 × 10^4^ vg/cell. Scatter plot of normalized gRNA A expression (y-axis) versus gRNA B expression (x-axis), each relative to U1 snRNA levels, for all tested promoter/terminator combinations. The blue inverted triangle indicates the top-performing combination (mU7v2.0 / HSUR4 for Cassette 1; RNU5B1 / RNU5A-8P for Cassette 2).

To identify higher-performing promoters, a panel of snRNA gene regulatory sequences **(Supplementary Fig. 5b)** was assessed via stable BxbI recombinase-mediated cassette integration **(see Methods)** in a GFP-G67R fluorescent reporter cell line **(Supplementary Fig. 4b)**. RNU5B1 emerged as the top-performing promoter, conferring approximately two-fold greater activity relative to mU7v2.0 **(Supplementary Fig. 5b)**. Terminator candidates were subsequently identified by screening a curated library of 533 human snRNA expression cassettes **(see Methods)** in the same reporter line, from which RNU5A-8P and RNVU1-31 were selected as lead terminator candidates **(Supplementary Fig. 5d–e)**. In parallel, snRNA homologues derived from Herpesvirus saimiri U-RNA elements (HSURs) were evaluated on the basis that viral regulatory sequences are subject to evolutionary pressure favoring maximal transcriptional output^26^. Among those tested, HSUR1, HSUR3, and HSUR4 conferred equivalent or modestly superior reporter expression relative to RNU5B1 **(Supplementary Fig. 5f)**, leading to selection of the HSUR4 terminator for incorporation into the final cassette design.

Having established high-performance regulatory elements, we next determined the optimal number of expression cassettes per AAV vector. Given the compact footprint (~700 bp) of each cassette, we employed a self-complementary AAV (scAAV) vector platform, which bypasses host-cell second-strand synthesis to achieve faster expression kinetics and enhanced potency relative to single-stranded AAV (ssAAV)^27^. Within the scAAV platform, we sought to maximize payload expression by incorporating multiple cassettes while minimizing recombination driven by inter-cassette sequence homology. We found that vectors encoding three or four size-reduced cassettes produced highly heterogeneous viral populations incompatible with therapeutic use due to sequence homology driven recombination. Single and dualcassette vectors demonstrated minimal heterogeneity, remaining over 80% or 70% intact, respectively, across multiple vector lots **(Supplementary Fig. 6a–b)**. Single- and dual-cassette vectors **(Fig. 2b)** were subsequently evaluated in a dose-titration experiment, with the dual-cassette configuration demonstrating significantly greater on-target editing efficiency at the highest doses **(Fig. 2c)**, supporting advancement of the two-cassette architecture to promote higher gRNA expression and activity from a single AAV genome.

Finally, lead promoter and terminator elements were assembled combinatorially with two unique tool gRNAs, packaged into AAV, and transduced into ARPE-19 cells. gRNA expression was quantified by droplet digital PCR (ddPCR) and normalized to endogenous U1 snRNA. The highest-performing con-figuration for both gRNAs paired the mU7v2.0 and RNU5B1 promoters with the HSUR4 and RNU5A-8P terminators, respectively **(Fig. 2d)**. This architecture was designated the final vector design for packaging MECP2-02, MECP2-03, and MECP2-04 gRNAs for delivery to disease-relevant model systems.

### Precision RNA editing in iPSC-derived neurons restores MeCP2 protein expression

To evaluate the therapeutic potential of ADAR-mediated RNA editing for the correction of the R168X nonsense mutation, we first assessed the functionality of optimized snRNA-gRNA payloads in neurons derived from iPSCs generated from RTT patients harboring the *MECP2*^*R168X*^ mutation. The two isogenic lines used for experiments were commercially procured as two lines from the same donor cell pool that had been identified to express the R168X allele (R168X X^A^) or the WT allele (WT X^A^). However, through RNA sequencing of differentiated neurons from each line, we observed that while 96% of total MeCP2 transcripts in line R168X X^A^ displayed the R168X allele, 36% of transcripts in WT X^A^ cells also represented R168X **(Supplementary Fig. 7)**, indicating a drift in X-inactivation status or leaky expression of the R168X allele in neurons derived from the WT X^A^ line. Both lines were treated with the three different gRNAs packaged into the optimized snRNA expression framework (**Fig. 3a)** using AAV-PhP.eB at either 5 × 10^4^ vg/cell (low dose) or 5 × 10^5^ vg/cell (high dose). Following 7 days in culture, we evaluated editing at the target adenosine using Sanger sequencing. MECP2-02 led to >20% editing in line R168X X^A^ at both doses, but yielded significantly lower editing in WT X^A^ neurons **(Fig. 3b)**, indicating conservation of allele specificity of the MECP2-02 gRNA in patient-derived neurons. MECP2-03 and MECP2-04 led to significantly higher levels of RNA editing, with more than 50% across both lines, and did not differ significantly across the two lines at the highest dose **(Fig. 3b)**. To characterize the local sequence specificity of ADAR recruitment and to identify any proximal offtarget editing events, we further analyzed Sanger sequences containing the entire gRNA hybridization region flanking the target adenosine. Chromatogram analysis revealed a clean editing signature restricted to the target adenosine, with no evidence of bystander editing at neighboring adenosines within the hybridization domain, underscoring the precision of the snRNA-gRNA architecture in human iPSC derived neurons **(Fig. 3c)**. Restoration of full-length MeCP2 protein was confirmed via Western blot in line R168X X^A^, with cells treated with high dose MECP2-03 and MECP2-04 showing a clear increase in MeCP2 protein compared to untreated or control treated cells **(Fig. 3d)**. Together, these findings demonstrate that snRNA-guided ADAR-mediated RNA editing can efficiently and precisely correct the MeCP2 R168X non-sense mutation in patient-derived neurons, restoring full-length MeCP2. Notably, the absence of detectable bystander editing highlights the precision of this approach, supporting its translational potential as a therapeutic strategy for Rett syndrome.

**Fig. 3:**
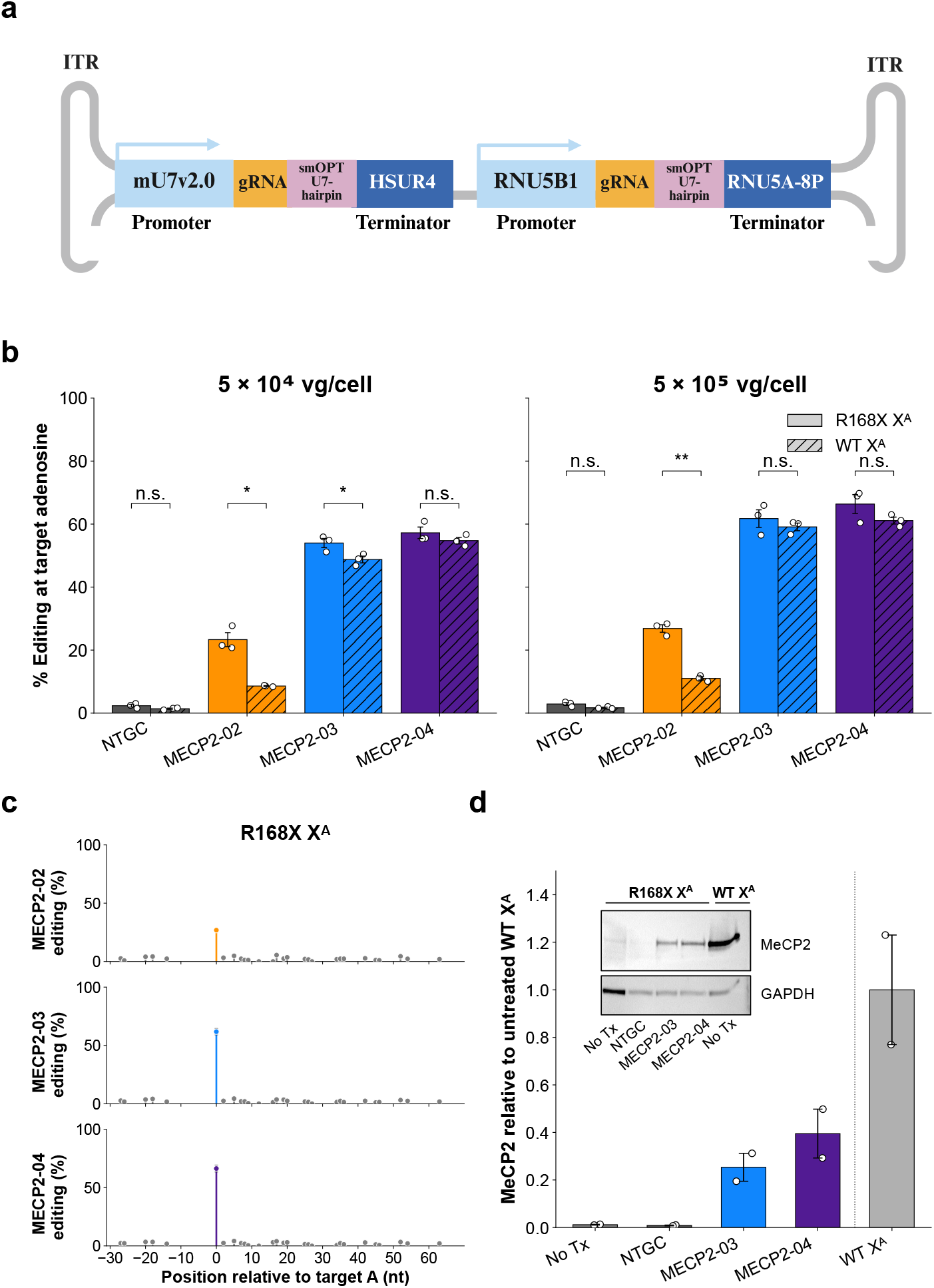
AAV-snRNA-gRNA correction of the R168X mutation and restoration of protein expression in RTT patient iPSC-derived neurons. **a**, Schematic of the dual-guide AAV construct used for programmable ADAR-based RNA editing. The vector, flanked by inverted terminal repeats (ITRs), encodes two expression cassettes: one driven by the mU7v2.0 promoter expressing a gRNA with a previously optimized hairpin structure and under the control of the HSUR4 terminator, and a second driven by the RNU5B1 promoter expressing a second identical gRNA under the control of the RNU5A-8P terminator. Created in BioRender. Vanschoiack, A. (2026) https://BioRender.com/9rt70z8. **b**, Percent editing at the target adenosine in R168X X^A^ (solid bars) and WT X^A^ (hatched bars) hiPSC-derived neurons treated with three gRNA vectors (MECP2-02, MECP2-03, MECP2-04) or a non-targeting guide control (NTGC) at two AAV doses: 5×10^4^ vg/cell (left) and 5×10^5^ vg/cell (right). Statistical comparisons between R168X X^A^ and WT X^A^ are indicated (Two-sided Welch’s t-test, n.s. = not significant, * p < 0.05, ** p < 0.01). Data points represent replicate experiments; bars show mean ± SEM. **c**, Local off-target editing analysis for R168X X^A^ cells treated with MECP2-02 (top), MECP2-03 (middle), and MECP2-04 (bottom) at 5×10^5^ vg/cell. Editing frequency (%) is plotted at all adenosine positions relative to the target adenosine (position 0). Colored bars at position 0 highlight on-target editing; surrounding positions reflect minimal bystander events. **d**, MeCP2 protein quantification and representative western blot from R168X X^A^ cells treated at 5×10^5^ vg/cell. Lower: MeCP2 levels normalized to the WT X^A^ no-treatment control (WT–No Tx) for untreated R168X cells (No Tx), NTGC, MECP2-03, and MECP2-04 conditions. Data shown as mean ± SEM. Upper: Western blot probed for MeCP2 and GAPDH (loading control) across R168X (No Tx, NTGC, MECP2-03, MECP2-04) and WT X^A^ (No Tx) conditions.

### Transcriptome-wide profiling of AAV-snRNA-gRNA expression in iPSC derived neurons

MECP2-03 and MECP2-04 gRNAs were further evaluated by RNAseq to assess transcriptome-wide specificity of the top performing gRNAs. RNAseq evaluation of on-target A-to-I editing confirmed slightly increased levels of dose-dependent editing on both R168X and WT MeCP2 alleles compared to Sanger evaluation. At the high dose (5 × 10^5^ vg/cell), MECP2-03 and MECP2-04 achieved 85.3 ± 4.7% and 83.5 ± 11.1% editing in R168X X^A^ neurons, respectively, and 80.6 ± 3.5% and 81.7 ± 5.0% in line WT X^A^. At the low dose (5 × 10^4^ vg/cell), the observed editing was 62.5 ± 4.7% and 69.2 ± 8.8% in R168X X^A^ neurons, and 53.6 ± 15.6% and 50.9 ± 14.0% in WT X^A^ neurons. No editing was detected in the non-targeting gRNA (NTGC) or vehicle controls. Bystander A-to-I editing was assessed at 29 adenosine positions within the gRNA hybridization window. Mean bystander editing did not exceed 2% at any position, compared to 0.5% background in NTGC and vehicle controls **(Fig. 4a)**.

**Fig. 4:**
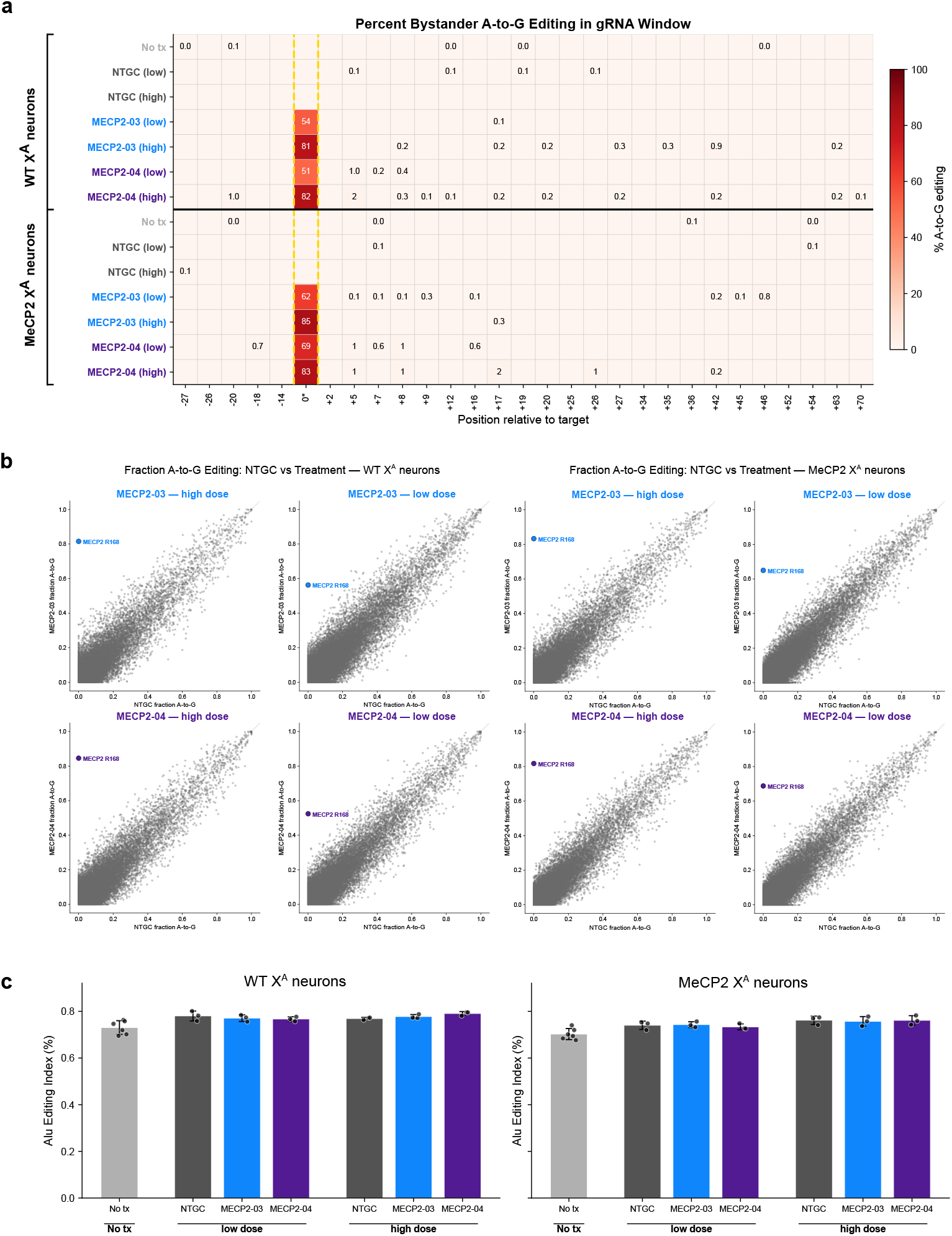
Transcriptome-wide specificity of lead AAV-Sn-gRNA constructs, in iPSCs. **a**, Bystander editing heatmap showing mean A-to-I editing rates across all adenosines within the gRNA hybridization region (positions relative to the target site) for each lead gRNA payload, stratified by cell type (WT and R168X). High editing at the target adenosine (position 0) with minimal off-target activity within the window is shown. **b**, Transcriptome-wide off-target A-to-I editing scatter plots generated from JACUSA2 analysis, comparing edited samples against NTGC control. Each point represents an editome site; the x-axis shows editing fraction in the NTGC control and the y-axis shows editing fraction in the gRNA-treated sample. Points above the diagonal indicate gRNA-dependent off-target editing. Left panel: WT allele samples; right panel: R168X allele samples. The on-target edit is highlighted in light blue (MECP2-03) or purple (MECP2-04). **c**, Global ADAR editing quantified by the Alu editing index (AEI) for each treatment group and genotype. Each bar represents the mean AEI for each sample with individual replicates as black circles and error bars represent the standard deviation.

Differential expression analysis confirmed that *MECP2* transcript levels were unchanged between payload-treated and NTGC-treated neurons from both iPSC lines (DESeq2; all padj > 0.99), indicating that neither the synonymous edit in WT cells (CGA→CGG) nor the corrective edit in R168X cells (TGA→TGG, X168W) alters mRNA stability. *MECP2* expression was approximately 2-fold lower in neurons derived from line R168X X^A^ relative to WT X^A^ (log_2_FC = ™0.90, padj = 1.5 × 10^™62^), potentially reflecting differences in nonsense mediated decay due to R168X mutant transcript or clonal variation during cell line selection.

To identify potential off-target hybridization sites, BLAST alignment of the MECP2-03 and MECP2-04 gRNA sequences was performed against the human genome **(see Methods)**. This identified 139 putative hybridization sites for each payload, mapping to 134 and 136 annotated genes for MECP2-03 and MECP2-04, respectively. The MECP2-03 and MECP2-04 gRNAs share 93% sequence identity (95/102 nt), differing only in a 13-nt central window; accordingly, 94 of these genes were shared between the two payloads, with 40 unique to MECP2-03 and 42 unique to MECP2-04.

To directly evaluate whether gRNA hybridization at these predicted sites produces measurable tran-scriptomic effects, we performed a focused analysis of the BLAST hit gene panel to screen for hybridization dependent off-target editing, splicing, or gene expression changes. 97/134 MECP2-03 and 98/136 MECP2-04 BLAST hit genes were detectably expressed in neurons derived from iPSC lines R168X X^A^ and WT X^A^ (mean TPM ≥ 0.1). At the gene expression level, no BLAST hit genes were differentially expressed in any payload-versus-NTGC contrast (DESeq2; padj < 0.05, |log_2_FC| > 1.0). Similarly, differential splicing analysis restricted to BLAST hit genes identified no significant events in any payload-versus-NTGC comparison (rMATS; FDR < 0.05, |ΔPSI| ≥ 0.25 or ≥ 0.10 for near-constitutive exons, junction read depth ≥ 100). Additionally, no evidence of off-target A-to-I editing was observed within BLAST hit sites (JACUSA log-likelihood ratio > 10, A-to-G change > 0.05, read depth > 50). This confirms that, despite partial sequence complementarity at these loci, gRNA hybridization does not produce detectable changes in gene expression, splicing, or ADAR mediated editing at any predicted off-target site (**Table S1**).

Having confirmed no detectable effects at predicted hybridization sites, we next assessed the full transcriptome for payload-induced changes (i.e., hybridization independent events). Site-level A-to-I editing fractions were highly concordant between payload-treated and NTGC-treated neurons, with the MeCP2 R168X target site as the sole clear outlier (**Fig. 4b**), qualitatively illustrating the absence of off-target editing. Formal differential editing analysis (JACUSA2; see Methods) confirmed this impression, identifying 168–328 hybridization-independent A-to-G events per contrast at the high dose, but these were classified as stochastic noise: 89–98% had JACUSA scores between 10 and 15 (just above the significance threshold) with median Δ editing of 0.11–0.12, site-level cross-genotype reproducibility was near zero (0–3 shared sites per payload), and cross-payload concordance was similarly negligible (6–9 shared sites) despite 93% gRNA sequence identity. Zero sites were shared across all four high-dose contrasts; at the low dose, only 0–2 events were detected. Global ADAR function was also preserved: the Alu Editing Index (AEI) confirmed there was no appreciable difference in global Alu editing between any gRNA treatment and the dose-matched NTGC control in both genotypes (two one-sided tests, equivalence margin ±0.05 percentage points AEI; all p ≤ 0.039). (**Fig. 4c**).

Differential expression analysis (PyDESeq2; see Methods) identified no differentially expressed genes (DEGs) for MECP2-03 at any dose or cell line. MECP2-04 yielded 6 DEGs in WT X^A^ and 7 in R168X X^A^ at the high dose, with only one gene (HMGB2) shared between genotypes; no DEGs were detected at the low dose for either payload. The minimal cross-genotype overlap (1 of 12 unique genes), absence of dose-responsiveness, and lack of cross-payload concordance are consistent with stochastic variation rather than systematic off-target activity.

Global differential splicing analysis (rMATS; see Methods) identified 17–81 hybridization-independent events per contrast at the high dose. At the low dose, events were nearly eliminated (0–3 per contrast). Cross-genotype concordance was low (3 of 141 MECP2-03 events; 0 of 63 MECP2-04 events), and cross-payload overlap was similarly limited (10 shared genes out of >150 unique), despite the two gRNAs sharing 93% sequence identity and 94 of their 134–136 BLAST-predicted target genes. The majority of events had |ΔPSI| values near the detection threshold (0.10–0.12), consistent with stochastic variation.

MECP2-03 and MECP2-04 gRNAs demonstrated a favorable specificity profile across all three offtarget mechanisms evaluated: differential expression, alternative splicing, and A-to-I editing. Focused analysis of 134–136 genes at BLAST-predicted gRNA hybridization sites revealed no differential expression, splicing, or editing changes in any payloadversus-NTGC comparison. Transcriptome-wide, zero hybridization-dependent off-target events were detected in any modality, dose, or genotype (**Table S1**). The hybridization-independent expression, splicing, and editing changes observed at the high dose showed minimal reproducibility across cell lines and payloads, were not dose-responsive, and are consistent with stochastic variation at the respective statistical thresholds rather than systematic off-target activity. Importantly, global ADAR function was preserved despite recruitment of endogenous enzymes to a non-native target sequence. These results support the transcriptome-wide specificity of MECP2-03 and MECP2-04 gRNAs when applied to the MeCP2 R168X target.

### Lifespan extension in RTT mice via single AAV-snRNA-gRNA administration

To assess the therapeutic efficacy of snRNA-gRNAs *in vivo*, all three gRNA payloads: MECP2-02, MECP2-03, and MECP2-04, were packaged into the CNS-tropic AAV-PHP.eB capsid and administered to *Mecp2*^*R168X*^ male mice intravenously (**Fig. 5a)**. All treatment cohorts received a single retro-orbital (RO) injection between postnatal days 14–19 at a dose of 5 × 10^13^ vg/kg, alongside a vehicle-injected control group (n = 10 per group). Cohort 1 (n = 5 per group) was sacrificed at four weeks post-treatment to quantify Mecp2 RNA editing efficiency and the degree of protein restoration in the brain. Cohort 2 animals were monitored longitudinally for up to 184 days post-injection, with serial assessments of body weight, Rett Phenotype Score^28^, and survival.

**Fig. 5:**
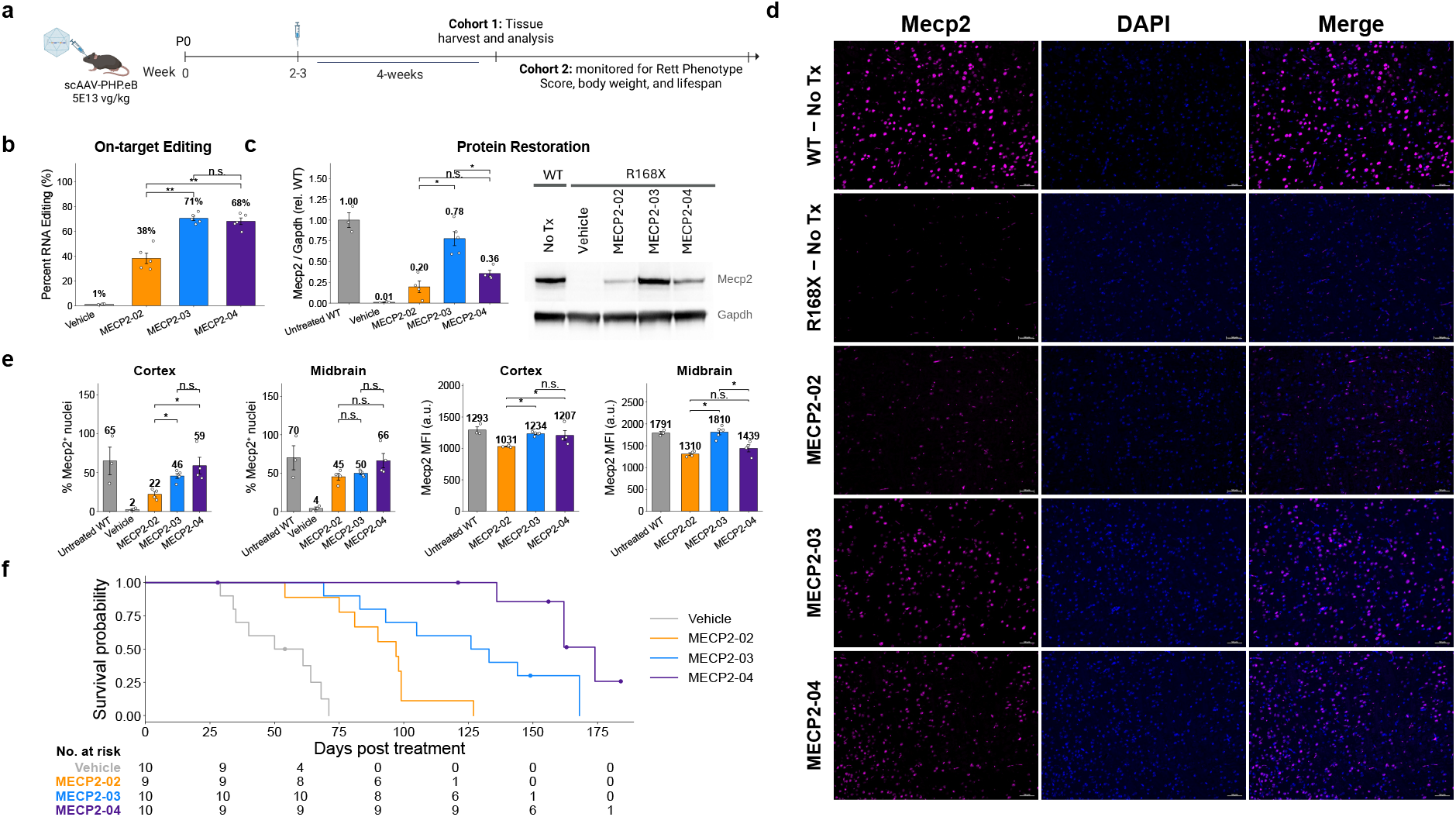
Programmable ADAR-based RNA editing extends survival in a mouse model of Rett syndrome. **a**, Experimental schematic. Mice received a single intravenous injection of scAAV-PHPeB (5×10^13^ vg/kg) at postnatal days 14 - 19. Cohort 1 was sacrificed at 4 weeks post-injection for tissue harvest and molecular analyses; Cohort 2 was monitored longitudinally for Rett phenotype score, body weight, and lifespan. Created in BioRender. Vanschoiack, A. (2026) https://BioRender.com/mza2wia. **b**, On-target RNA editing efficiency. Bar graph showing percent RNA editing at the target adenosine in mice treated with vehicle or three guide RNA constructs (MECP2-02, MECP2-03, MECP2-04). **c**, Mecp2 protein restoration. (Left) Quantification of Mecp2 protein levels normalized to Gapdh relative to untreated wild-type (WT) controls. (Right) Representative western blot showing Mecp2 and Gapdh bands across Cohort 1 treatment groups in R168X mutant mice and WT controls. **d**, Immunofluorescence imaging of Mecp2 protein expression in brain sections. Columns show Mecp2 signal (magenta), DAPI nuclear stain (blue), and merged images across five conditions: WT–No Treatment, R168X–No Treatment, and R168X mice treated with MECP2-02, MECP2-03, or MECP2-04. Scale bars = 50µm. **e**, Quantification of Mecp2 immunofluorescence in cortex and midbrain. (Left panels) Percentage of Mecp2-positive nuclei in the cortex and midbrain across treatment groups. (Right panels) Mean fluorescence intensity (MFI) of Mecp2 signal (arbitrary units) in cortex and midbrain. Statistics for all bar charts (b, c, e): two-sided Mann-Whitney U tests on raw (uncorrected) p-values; \**p* < 0.05, \*\**p* < 0.01, \*\*\**p* < 0.001, n.s. = not significant. Bars represent mean ± SEM. **f**, Kaplan-Meier survival analysis. Survival curves for Vehicle (gray), MECP2-02 (orange), MECP2-03 (blue), and MECP2-04 (purple) treated R168X mice over 175 days post-treatment (n = 9–10 pergroup). Numbers at risk are shown below the x-axis.

To determine whether systemic AAV delivery achieved sufficient CNS transduction, on-target editing activity, and restoration of Mecp2 protein the left cerebral hemisphere (excluding the olfactory bulbs and cerebellum) was dissected for RNA and protein extraction from lysate, while the right hemisphere was processed for histological assessment of Mecp2 protein restoration from cohort 1 animals. Targeted RT-PCR and Sanger sequencing confirmed robust ontarget editing across all mice administered Mecp2-targeting payloads. Among the constructs evaluated, MECP2-03 achieved the highest mean editing efficiency at 70.6 ± 4.1%, followed by MECP2-04 at 68.2 ± 6.1% and MECP2-02 at 38.3 ± 9.5% (**Fig. 5b**). Notably, these editing frequencies were derived from wholebrain tissue, suggesting widespread viral transduction and editing activity throughout the CNS. Western blots from brain lysates confirmed detection of full-length Mecp2 protein with all three treatments, consistent with RNA editing re-establishing the fulllength open reading frame, with MECP2-03 leading to significantly higher restoration of bulk protein compared to MECP2-02 and MECP2-04 (p < 0.05) (**Fig. 5c**). Immunofluorescence evaluation across whole sagittal brain sections qualitatively demonstrated broad Mecp2 protein restoration throughout the brain with all three treatments (data not shown). Restored Mecp2 protein was qualitatively indistinguishable from wild-type Mecp2, following the expected nuclear localization with accentuated staining within heterochromatic foci (**Fig. 5d)**. Detailed analysis of percent of nuclei expressing full-length Mecp2 in cortical and midbrain regions revealed that the percent of cells with restored Mecp2 protein ranged from 22.1% (cortex) to 45.2% (midbrain) for MECP2-02, 45.7% (cortex) and 49.7% (midbrain) for MECP2-03, and 58.9% (cortex) to 64.2% (midbrain) for MECP2-04 (**Fig. 5e**). Further evaluation of the mean fluorescence intensity (MFI) of the Mecp2 positive nuclei revealed that MECP2-03 led to significantly higher MFIs compared to MECP2-02 in cortex (p < 0.05) and both MECP2-02 and MECP2-04 in midbrain (p < 0.05), consistent with higher total protein with MECP2-03 as detected by Western blot, indicating that MECP2-03 is most efficient at restoring protein in transduced cells.

Animals from the survival cohort were assessed using a composite Rett phenotype scale encompassing mobility, gait, hindlimb clasping, tremor, and general condition, where higher scores reflect greater disease severity^28^. Given the rapid attrition of vehicle-treated animals, which confounds between-group comparisons at later time points, longitudinal analyses were restricted to a data-driven observation window spanning days 0–43 post-treatment, defined as the final assessment day on which at least five vehicle-treated animals contributed phenotype data. Across this window, all three treatment groups exhibited attenuated phenotype progression relative to vehicle. Linear mixed-effects (LME) models incorporating random intercepts and random time slopes per animal identified significant treatment-by-time interactions for the aggregate phenotype score: MECP2-03 (β = ™0.093, p < 0.001), MECP2-02 (β = ™0.078, p < 0.001), and MECP2-04 (β = ™0.076, p < 0.001), indicating that all three treatments significantly reduced the daily rate of phenotype worsening relative to vehicle **(Supplementary Fig. 8)**. At the sub-score level, mobility showed the most consistent treatment effect, with all three payloads significantly slowing decline (p < 0.02 for each). Tremor scores were reduced in the MECP2-02 (p = 0.04) and MECP2-03 (p = 0.08) groups. Gait and general condition sub-scores showed directionally favorable but non-significant trends across all treatment groups, while hindlimb clasping was minimal in all groups, likely reflecting limited statistical power within the 43-day observation window. Body weight trajectories were largely comparable across groups throughout this period, although MECP2-03 showed modest preservation relative to vehicle (p = 0.02).

Beyond slowing early phenotypic deterioration, treatment with all three constructs also produced a pronounced and statistically robust extension of survival. Notably, treatment with all three gRNA payloads conferred marked survival extensions relative to vehicle-treated controls (**Fig. 5f**). Animals euthanized for intercurrent, non-Rett-related causes (e.g., severe tail lesion, malocclusion; n = 4) and those surviving to study termination (n = 5) were right-censored in the survival analysis. Vehicle-treated *Mecp2*^*R168X*^ mice exhibited a median survival of 50 days, consistent with the expected disease trajectory^29^. By contrast, median survival increased to 97, 126, and 174 days for MECP2-02, MECP2-03, and MECP2-04, respectively, representing a 3.5-fold extension for the best-performing construct. Pairwise log-rank tests confirmed highly significant differences in survival for all treatment groups relative to vehicle (MECP2-02: p = 0.0002; MECP2-03: p < 0.0001; MECP2-04: p < 0.0001; Holm-corrected). Penalized Cox proportional hazards regression yielded hazard ratios of 0.12 (95% CI: 0.03–0.49), 0.09 (95% CI: 0.02–0.37), and 0.07 (95% CI: 0.02–0.35) for MECP2-02, MECP2-03, and MECP2-04, respectively, corresponding to an 88–93% reduction in the instantaneous risk of death. Complementary restricted mean survival time (RMST) analysis, which does not assume proportional hazards, corroborated these findings, with treated animals gaining an estimated 39, 74, and 115 additional days of survival relative to vehicle for MECP2-02, MECP2-03, and MECP2-04, respectively, over the 184-day observation period (all p < 0.001). The results are summarized in (**Supplementary Fig. 8c-d)**. Notably, three MECP2-04 and two MECP2-03 treated animals remained alive at study termination and were right-censored, suggesting the true survival benefit for these cohorts may be modestly underestimated.

To contextualize these inter-construct differences in survival, pairwise comparisons were conducted across both molecular and phenotypic endpoints to identify the factors most likely to underlie the observed hierarchy of therapeutic benefit. Both MECP2-03 and MECP2-04 gRNAs conferred significantly greater survival benefit than MECP2-02 (all pairwise Holm-corrected log-rank p ≤ 0.031), a result consistent with the significantly lower editing efficiency observed for MECP2-02 (38.3%, p < 0.05). Importantly, despite achieving comparable RNA editing levels (70.6% vs. 68.2%; p = 0.69), MECP2-04 con-ferred significantly greater survival than MECP2-03 (median 174 vs. 126 days; Holm-corrected log-rank p = 0.031; RMST difference = 41 days, p = 0.003). Longitudinal mixed-effects modeling of all phenotypic sub-scores revealed no significant differences in the rate of early phenotype progression among any treatment group pair during the 43-day analysis window (all Holm-corrected treatment-by-time interaction p > 0.13), indicating broadly comparable early disease trajectories across payloads. The dissociation between equivalent early phenotypic trajectories and divergent long-term survival outcomes for MECP2-03 versus MECP2-04 suggests that the survival advantage conferred by MECP2-04 is mediated by factors not captured by early phenotype scoring. Evaluation of bulk brain editing from the limited animals euthanized at termination of the survival study revealed higher editing with MECP2-04 (62.1% and 60.2% editing at end of 184-day ob-servation period) compared to MECP2-03 (39.6% and 52.8% editing at end of 184-day observation period) (**Supplementary Fig. 8d)**, though more animals would need to be evaluated at later timepoints to determine whether this is a treatment-related effect and correlated with improved survival. Complete pairwise comparisons across all endpoints are provided in Table S2. Taken together, these findings demonstrate that AAV-PHP.eB-mediated delivery of snRNA-gRNA payloads targeting the *Mecp2*^*R168X*^ mutation achieves robust on-target RNA editing and widespread protein restoration, which collectively translate into significant attenuation of RTT-like phenotypes and a marked extension of survival in a severe hemizygous male mouse model. The graded therapeutic hierarchy observed across the three constructs with MECP2-04 outperforming MECP2-03 despite equivalent editing efficiency and lower total protein restoration underscores that editing efficiency alone is an incomplete predictor of *in vivo* therapeutic outcomes, and highlights the importance of comprehensive molecular profiling, including off-target analysis, in the prioritization of lead candidates for further preclinical and translational development.

## Discussion

This study establishes an integrated framework for precision therapeutic RNA editing that bridges generative AI design with the rational re-engineering of native RNA biogenesis pathways. Our results demonstrate that the synergy between DeepREAD-optimized gRNAs and enhanced snRNA expression scaffolds creates a robust, programmable platform capable of achieving systemic correction in the CNS that mitigates RTT phenotypes and extends lifespan in an RTT mouse model. By co-opting the cell’s own processing machinery to correct a pathogenic *MECP2* nonsense mutation, we bypassed the immunological and off-target risks associated with exogenous DNA editors, providing a modular blueprint for the treatment of a broad class of neurodevelopmental disorders.

We utilized our DeepREAD model to design highly efficient and specific gRNAs targeting the R168X codon. While the 5’-G flanking motif at the R168X locus is typically refractory to ADAR activity, our DeepREAD-directed generative design successfully navigated this context to achieve the high levels of editing required for functional rescue. Most notably, the model-driven approach went beyond mere potency optimization and generated a novel molecular “switch” within the MECP2-02 candidate that enables allelic discrimination. By utilizing an A at the −2 position, the gRNA enables a stable GA-GC bulge on the R168X transcript that stimulates editing, while collapsing into an inactive bulge against the WT allele.

While this highlights the capacity of generative AI to engineer allelic discrimination, we observed that this specific architecture was less potent in neurons compared to the allele cross-reactive candidates, MECP2-03 and -04. This suggests that for this target the structural constraints required for allelic specificity may inadvertently conflict with the catalytic requirements of ADAR2, the dominant isoform in the CNS. Consequently, the cross-reactive gRNAs emerged as the superior therapeutic lead, as they appear better optimized for the neuronal environment. Beyond their higher potency, these cross-reactive gRNAs offer distinct advantages for translational development: by directing a synonymous edit on the WT allele, they facilitate essential pre-clinical dosing and safety studies in non-human primates (NHPs), where no R168X disease model exists.

Previous attempts to edit Mecp2 RNA in an RTT mouse model required the co-delivery of exogenous ADAR2 and a gRNA in a single AAV genome, providing proof-of-concept; however, the off-target effects of supraphysiological levels of ADAR preclude its therapeutic use^7^. Complementing gRNA design, the optimization of the snRNA expression scaffold was essential for achieving therapeutic levels of editing. While this scaffold system was originally conceptualized in the 1990s for antisense-mediated exon skipping^30^, our previous work and current findings demonstrate that further improvements to the 5’ and 3’ regulatory elements can significantly enhance its ability to drive high and sustained expression. Beyond driving expression of gRNAs to recruit endogenous ADAR, these regulatory elements, promoters and terminators, could likely be readily applied to improve the efficiency of exon skipping, to express RNA aptamers, or even as a vector for gene replacement therapy in neurodevelopmental disorders caused by mutations in native snRNA genes^31^. Furthermore, our 2X tandem cassette architecture demonstrated a high degree of genomic stability within the AAV vector, a critical requirement for clinical-grade manufacturing; while also enabling multiplexed expression of gRNAs to alter multiple RNA targets^32^. This modularity establishes the snRNA scaffold as a versatile platform for delivering complex RNA payloads without the need for exogenous proteins.

The transcriptome-wide evaluation of MECP2-03 and MECP2-04 gRNAs revealed a highly favorable specificity profile, with no notable transcriptomic perturbations identified across our analysis. Because our platform co-opts endogenous cellular machineries by utilizing the snRNA pathway for gRNA biogenesis and endogenous ADAR for site-specific editing, it was critical to specifically interrogate both RNA splicing patterns and the global editome for off-target effects. Our analysis confirmed that these pathways re-mained functionally intact. Notably, while gene expression differences exist between untreated WT and R168X cells, the successful correction of *MECP2* did not immediately induce a large-scale shift in DEGs *in vitro*. A single gene was differentially expressed in both MECP2-03 and MECP2-04 treated cells compared to the NTGC negative control; however, this was not differentially expressed between WT and R168X cells. This likely reflects a timing issue inherent to the biological mechanism: the system requires a sufficient window for viral transduction, gRNA accumulation, and the subsequent synthesis and nuclear localization of full-length MeCP2 protein before downstream transcriptional targets can be remodeled. The fact that we observed profound phenotypic and survival improvements *in vivo* suggests that functional restoration occurs over a longer temporal window than a 7-day *in vitro* assay allows. Consequently, future *in vivo* studies that achieve measurable phenotypic rescue should incorporate large-scale transcriptomic profiling of the CNS to fully characterize the molecular restoration of MeCP2-regulated gene networks and the normalization of the disease-associated signature.

Our *in vivo* results suggest that a specific threshold of *MECP2* correction may be required to drive substantial extension of survival. While MECP2-02 provided a measurable benefit, its lower editing efficiency (~38%) was paired with a significantly shorter median survival compared to the MECP2-03 and MECP2-04 cohorts, which both achieved ~70% editing. However, average editing efficiency did not explain all outcomes; notably, MECP2-04 conferred a statistically significant survival advantage over MECP2-03 (174 vs. 126 days) despite achieving nearly identical bulk editing levels and significantly higher bulk protein detected with MECP2-03 compared to MECP2-04. This divergence implies that the long-term therapeutic outcome may be influenced by factors beyond aggregate efficiency, such as the precision of the gRNA’s specificity profile or the degree of correction within critical, albeit small, neuronal sub-populations that disproportionately impact survival. Administration of MECP2-03 and -04 to WT mice could further inform if off-target activity or adverse events contribute to the lower efficacy of MECP2-03.

The timing and delivery of this postnatal intervention is an important consideration for clinical translation. Our treatment window (P14–19) roughly parallels the 12–24 month developmental stage in humans^33^, which typically coincides with the onset of clinical regression in RTT^34^. This indicates that RNA editing could serve as a viable interventional strategy even after symptomatic onset, potentially arresting or reversing disease progression. Regarding delivery, while AAV-PHP.eB provided the necessary CNS biodistribution in this study^8^, the translation to human patients will necessitate alternative strategies. Although we and others have reported novel AAV variants that can cross the BBB in NHP^35–40^, the field awaits clinical data to validate translation to humans. While systemically delivered brain-targeting AAV variants are being pursued, intracerebroventricular (ICV) or intrathecal (IT) routes remain established clinical alternatives. Meanwhile, advanced pre-natal genetic screening may allow for early interventions within the first year of life^41^, a developmental period when the BBB is permissive to wild-type AAV9 and therapeutic intervention may be more impactful^42^. These findings collectively support the feasibility of a single-administration to achieve durable therapeutic gains in RTT patients.

The success in identifying potent gRNAs for the R168X mutation provides a blueprint for targeting other common RTT-associated nonsense variants, such as R255X, R270X, and R294X. Our platform is compatible with the FDA’s newly released Plausible Mechanism Framework^43^, which allows for accelerated clinical expansion when multiple mutations are addressed using the same underlying technology and mechanism of action. By integrating precision engineering with this regulatory-friendly framework, we can rapidly expand the treatable patient population for RTT and other G-to-A genetic disorders.

## Methods

### Reporter Construct Design and Cloning

For the Human MeCP2 construct, a minigene encoding the wildtype human MeCP2 e1 isoform (NM_001110792.2) was synthesized by Twist Bioscience (South San Francisco, CA, USA) and supplied in a pTwist-CMV-WPRE-Neo backbone. The minigene was designed to contain a single canonical ATG start codon; all down-stream ATG codons within the open reading frame (ORF) were replaced with GTG to preclude alternative translation initiation. To facilitate synthesis, the polyalanine stretch at the N-terminus and the region encoding residues E386–P417 were codon-optimized. The R168X nonsense mutation (CGA>TGA) was subsequently introduced into the wild-type minigene by PCR-based mutagenesis using mutagenic primers and PrimeSTAR GXL DNA polymerase (Takara Bio, San Jose, CA, USA). Both the wild-type and R168X mutant MeCP2 minigene fragments were subcloned into a PiggyBac transposon expression vector, positioned between an EF1α promoter and an SV40 polyadenylation signal, using HiFi DNA Assembly (New England Biolabs [NEB], Ipswich, MA, USA) at a 3:1 molar insert-to-vector ratio. Prior to assembly, the recipient vector was linearized by double di-gestion with KpnI and BamHI (NEB), dephosphorylated with Quick CIP (NEB), and gel-purified to eliminate residual undigested vector. A FLAG epitope tag (DYKDDDDK) was incorporated in-frame at the C-terminus of each construct, immediately upstream of the stop codon, yielding the wt-MeCP2-FLAG and R168X-MeCP2-FLAG fusion reporter constructs.

Following assembly, 2 µL of each reaction was transformed into NEB Stable chemically competent *Escherichia coli* (NEB), and transformants were selected on LB agar supplemented with carbenicillin (LB-Carb; Teknova, Hollister, CA, USA). Individual colonies were picked and cultured overnight in LB broth containing 100 µg/mL carbenicillin. Plasmid DNA was isolated using the NucleoSpin Plasmid Purification Kit (endotoxin-free; Macherey-Nagel Inc., Allentown, PA, USA) and sequence-verified by Sanger sequencing prior to use.

For the Mouse Mecp2 construct, the native cDNA ORF corresponding to the wild-type mouse *Mecp2* e2 isoform (NM_010788.4) was synthesized by Genscript (Piscataway, NJ, USA) and supplied in a pcDNA3.1+/C-DYK backbone encoding a C-terminal FLAG tag (DYKDDDDK), analogous to the human reporter constructs described above. The R168X non-sense mutation (AGA>TGA) was introduced into the wild-type sequence by PCR mutagenesis using mutagenic primers and PrimeSTAR GXL DNA polymerase (Takara Bio). Both wild-type and R168X mutant mouse *Mecp2* cDNA fragments were subcloned into the same PiggyBac expression vector used for the human constructs, between the EF1α promoter and SV40 polyadenylation signal. Cloning, bacterial transformation, and plasmid preparation were carried out as described above.

### Generation of HEK293T-MeCP2-4iso Reporter cell line

HEK293 cells (ATCC, Manassas, VA, USA) were seeded at 1 × 10^5^ cells per well in CellBIND 24-well plates (Corning, Corning, NY, USA) and left undisturbed at 37°C for 6h to allow cell attachment prior to transfection. Transfections were performed using TransIT-293 transfection reagent (Mirus Bio, Madison, WI, USA) in Opti-MEM I Reduced Serum Medium (Gibco, Thermo Fisher Scientific, Waltham, MA, USA). For each transfection, 750 ng of reporter or transposon plasmid(s) and 250 ng of piggyBac transposase plasmid were combined with Opti-MEM to a total volume of 8 µL, followed by addition of 2 µL TransIT-293 (2:1 reagent-to-DNA ratio, v/w). Complexes were incubated at room temperature for 15 min, supplemented with 20 µL culture medium, and 30 µL of the final transfection mixture was added dropwise to each well before returning cells to a humidified 37°C, 5% CO_2_ incubator.

The pooled reporter cell line was established by co-transfecting all four MeCP2-Flag reporter constructs simultaneously, i.e. human wild-type MeCP2-Flag, human R168X MeCP2-Flag, mouse wild-type MeCP2-Flag, and mouse R168X Mecp2-Flag (750 ng total, equimolar amounts), together with 250 ng of the piggyBac transposase plasmid, enabling stable genomic integration of all four reporters within a single pooled population. All the transposon plasmids had a CMV-PuroR-bGH polyA cassette in addition to the MeCP2 reporter cassette, to enable antibiotic selection of cells with stable integration.

Seventy-two hours post-transfection, cells with stable integration of the reporters were selected by addition of puromycin (Life Technologies, Thermo Fisher Scientific, Waltham, MA, USA) to the culture medium at a final concentration of 5 µg/mL. Selection was maintained for 14 days, with complete cell death of non-transfected control wells used as the endpoint criterion for selection stringency. Surviving pooled populations were expanded and maintained in Dulbecco’s Modified Eagle Medium (DMEM; Gibco, Thermo Fisher Scientific) supplemented with 10% heat-inactivated fetal bovine serum (FBS; Gibco, Thermo Fisher Scientific) at 37°C, 5% CO_2_.

Stable integration was validated by flow cytometry. Functional assessment of the MeCP2-R168X reporters was performed by transient transfection of gRNAs targeting the R168X MeCP2 transcript, followed by total RNA isolation, reverse transcription, PCR amplification of the target region, and Sanger sequencing to confirm editing outcomes.

### gRNA screening

The HEK293T-MeCP2-4iso cell was engineered using the PiggyBac transposon system to express the full-length MeCP2 open-reading frame with a C-terminal Flag-tag for four variants: human MeCP2^R168X^, human MeCP2^WT^, mouse Mecp2^R168X^, and mouse Mecp2^WT^. gRNAs were cloned into a proprietary vector under the control of a mouse U7 snRNA promoter and 3′ regulatory elements. Engineered HEK293T cells were seeded 24 h before transfection in 96-well plates. gRNA transfections were performed using TransIT-293 Transfection Reagent (Mirus #2704) according to the manufacturer’s instructions. Cells were harvested 48 h post-transfection, and RNA was isolated using the RNeasy Mini Kit (Qiagen #74104). Reverse transcription was carried out with gene-specific primers and/or oligo dT utilizing SuperScript™ IV Reverse Transcriptase (Thermo Fisher #18090050). Amplicons were amplified utilizing KAPA HiFi DNA Polymerase (HotStart and Ready-Mix formulation) (Roche #07958935001). All samples were prepared for NGS with Illumina Nextera Unique primers and se-quenced on the Illumina iSeq.

### Sanger sequencing

*MeCP2* cDNA was amplified using Q5 High-Fidelity 2× Master Mix (NEB, #M0492) with primers 550F (5’-TTCGCTCTAAAGTAGAATTGATTGC-3’) and 769R (5’-TCTGATGCTGCTGCCTTT-3’), yielding a 220 bp amplicon. PCR cycling conditions were: 98°C for 30 seconds; 35 cycles of 98°C for 10 seconds, 64°C for 10 seconds, 72°C for 10 seconds; final extension at 72°C for 2 minutes. Products were verified by 2% E-gel electrophoresis and purified using the Zymo ZR-96 DNA Clean & Concentrator-5 kit (#D4024), eluting in 15 µL.

Purified PCR products were quantified by Qubit (dsDNA HS assay) and submitted in duplicate to Azenta Life Sciences for Sanger sequencing. Each well contained 10 ng of purified PCR product in 10 µL combined with 5 µL of 5 µM reverse primer for a total volume of 15 µL.

### RNA secondary structure generation

RNA duplex structures between gRNAs and *MeCP2* target sequences were predicted using the ViennaRNA package (v2.7.0) via its Python API. For each of gRNA–target combinations the RNA duplexfold was used to compute the minimum free energy structure and binding energy in dot-bracket notation. For visualization, the gRNA and target sequences were concatenated with a poly-A linker to form a hairpin substrate, and structure diagrams were rendered as SVG using VARNA (v3.93) with the naview layout algorithm. Base pairs were colored to distinguish Watson-Crick pairs from G-U wobbles, and the target adenosine edited by ADAR was highlighted.

### One copy and two copy snRNA vector screening

To assess whether there are improved outcomes with two RNAfix cassettes embedded within the AAV genome compared to one, two sets of constructs were cloned with one or two cassettes. The one cassette vectors contained the RNU5B1 promoter and RNU5A-8P terminator with synthetic filler sequence used to equalize the length of two-cassettes. The two cassette vectors had the first cassette as mU7v2.0 promoter and RNVU1-31 terminator and the second cassette contained the RNU5B1 promoter and RNU5A-8P terminator. In one iteration, the same gRNA was used in both cassettes and in the other iteration two different gRNAs, targeting the same adenosine, were used. In all iterations, the triple mutant mU7 hairpin was used as previously described^23^. These constructs were cloned, prepped to contain intact ITRs, and produced as AAV-DJ. These viruses were transduced onto WT HEK293 cells at MOIs of 5, 50, 250, 500, 2500, 5000, and 50,000. To assess editing of this model target, RNA was isolated at 48 hours post transduction. Protoscript II First Strand cDNA synthesis kit from NEB was used for cDNA which was then amplified by PCR and sequenced on an Illumina instrument to measure editing.

### Assessment of AAV Genomic Integrity

AAV genome integrity was evaluated by alkaline agarose gel electrophoresis. Prior to analysis, virus preparations were subjected to DNase I digestion (2 U per 1.5×10^11^ vector genomes [vg]) at 37°C for 60 minutes to eliminate residual unencapsidated nucleic acids. Encapsidated viral genomes were subsequently liberated and denatured by incubation in alkaline lysis buffer (50 mM EDTA, 1% SDS, 500 mM NaOH) for 20 minutes at ambient temperature. Denatured samples were electrophoretically resolved on a 1% alkaline agarose gel using 50 mM NaOH / 1 mM EDTA as the running buffer at a constant voltage of 100 V for 90 minutes. Following electrophoresis, gels were neutralized and stained with SYBR Gold nucleic acid stain for fluorescence-based visualization of genomic DNA species.

For orthogonal characterization of genome integrity, AAV genomic DNA was purified using the PureLink Viral RNA/DNA Minikit (Thermo Fisher Scientific; cat. no. 12280050) according to the manufacturer’s instructions. Purified DNA was submitted to Genewiz (Azenta Life Sciences) for long-read sequencing via PacBio Single Molecule, Real-Time (SMRT) sequencing technology. Resulting sequencing reads were aligned to a reference AAV genome sequence and annotated recombination product sequences to determine the percentage of full-length genomes within each preparation.

### SnRNA cassette screening

To select the final promoter and terminator choices of the two-cassette vector, the top performing promoters and terminators were arranged in different permutations and cloned in duplicate with two different gRNAs, in each instance both of the cassettes have the same gRNA. These vectors were then made into AAV and used to transduce ARPE-19 cells at an MOI of 1 × 10^4^ vg/cell. RNA was isolated at 48 hours post transduction, converted into cDNA with Protoscript II First Strand cDNA synthesis kit from NEB and gRNA expression was measured by ddPCR.

### Generation of an editing-dependent luciferase reporter

For screening purposes, a reporter construct was engineered to produce a luciferase signal as a quantifiable marker of deamination activity. To this end, a mini-gene harboring a target start codon along with its flanking 5’ untranslated region (UTR) and Kozak consensus sequence, followed by a downstream exon, was cloned upstream of a luciferase reporter in an alternative reading frame, separated by an intervening stop codon. Upon adenosine-to-inosine (A-to-I) deamination of the target start codon, translational initiation is redirected to the luciferase start codon, thereby shifting the reading frame and enabling luciferase expression.

### Generation of a GFP-G67R reporter by BxbI recombinase-mediated integration

To generate a cell line amenable to testing elements at a single copy per cell, we first inserted an attP1 site into the AAVS1 genomic locus (PPP1R12C) of HEK293T cells by CRISPR Cas9. Single cell clones were screened by PCR to obtain a population with one attP1 per cell. Next an intermediate cell line was created which contains an attB1 site, a cassette expressing GFP-G67R-P2A-mtagBFP to track both enrichment and GFP restoration, as well as a cassette driving blasticidin resistance and BxbI separated by another P2A with an attP2 in between the promoter and ORF. To enrich this intermediate cell line, blasticidin at a final concentration of 5 µg/mL was added and enrichment was assessed via BFP percent positive by flow cytometry. Prior the second integration enrichment was confirmed to be at least 90%. This second attP site can then be targeted by a plasmid containing an RNAfix cassette and an mCherry-P2A-Puromycin resistance cassette which only expresses upon targeted integration. The RNAfix cassette in this last plasmid can be modified to contain varying elements or a library of terminators for screening. Enrichment for the second integration is conducted by adding puromycin at 1.25 µg/mL final concentration.

### Flow-seq screen of snRNA terminators

A curated repertoire of 533 putative snRNA expression cassettes was assembled from RNAcentral and Ensembl and informed by published references^44^, with promoter regions defined as 300 bp upstream and terminator elements as 100 bp downstream of each transcription unit. The 100bp terminators were cloned via Golden Gate assembly in a shuttle vector downstream of three promoters; mU7v2.0, RNU5B1, RNU5E1 each paired with a constant gRNA and hairpin. The terminators were sequenced directly to confirm sufficient coverage across promoter replicates. The libraries were integrated in duplicate by plasmid transfection into the HEK293 GFP-G67R intermediate line with 100X coverage per variant with an integration efficiency assumed of 5%, so 20X the cell number was used. Controls of randomized DNA sequences, no gRNA cassette, were included for analysis references. The cells were expanded and maintained in high numbers to maintain diversity. The GFP intensities of RNU5B1 and mU7v2.0 promoter samples were similar enough to pool together for sorting. After achieving 90% for cells containing the cassette, the cells were sorted by GFP intensity using a Sony SH800S cell sorter with the top and bottom 10% isolated. At least ten million cells for each bin were acquired. The genomic DNA from those two bins and unsorted cells were isolated and sequenced on an Illumina Nextseq. Using a linear model of analysis, the terminators’ changes in abundance between the bins were assessed to identify those most enriched in the top 10% as a predictor of high performance.

### Cell Culture and AAV Transduction

P0 mixed mouse primary neurons were prepared from *Mecp2*^R168X/Y^ mice and WT littermates using standard methods^45^. Cells were cultured in Neurobasal medium with 25 μM glutamine, 1% penicillin–streptomycin, and B-27 supplement (Invitrogen) under a 5% CO2/10% O2 atmosphere at 37 °C. Neurons were transduced with MECP2-01 on D7 *in vitro* and maintained for seven days before fixing with 4% paraformaldehyde (PFA)/4% sucrose for immunostaining or harvesting in MagMAX lysis buffer containing 0.7% β-mercaptoethanol for RNA extraction.

RTT patient-derived iPSCs (System 1 Biosciences iPSC Line #125 clone 125.2/R168X X^A^ 125.9/WT X^A^) were engineered to express a Tet inducible NeuroD1 cassette using PiggyBac transposon system and differentiated into glutamatergic neurons using a doxycycline-inducible NeuroD1 rapid differentiation protocol modified from Zhang et al. (2013)^46^. Briefly, iPSCs were dissociated with Accutase and plated as single cells on Geltrex-coated 48-well plates in NeuroD1 Differentiation Medium 1 (DMEM/F12, N2 supplement, NEAA) supplemented with BDNF (10 ng/mL), NT-3 (10 ng/mL), doxycycline (1 μg/mL), laminin (0.2 μg/mL), and ROCK inhibitor (10 μM). ROCK inhibitor was removed after 24 hours. On day 2, cultures were transitioned to NeuroD1 Differentiation Medium 2 (Neurobasal, B27, GlutaMax, 2× Culture-One) with BDNF, NT-3, and doxycycline. On day 6, a 50% media change was performed with 0.25 μM Ara-C and Geltrex (1:100) to suppress proliferating non-neuronal cells, followed by a complete media change on day 7 to remove Ara-C. On day 8 of differentiation, neurons were transduced with self-complementary AAV vectors encoding MECP2-03 and MECP2-04 gRNAs, or a non-targeting gRNA control (NTGC) at two doses (5 × 10^4^ or 5 × 10^5^ vg/cell); untransduced wells served as Vehicle controls. Cultures were maintained for 7 days post-transduction with media changes every 2–3 days. Cells were lysed in RIPA buffer (Thermo Fisher, #89901) supplemented with HALT Protease and Phosphatase Inhibitor Cocktail (1:100; Thermo Fisher, #78441) for Western blot anal-ysis, MagMAX lysis/binding buffer containing 0.7% β-mercaptoethanol, or fixed in 4% PFA/4% sucrose for immunostaining.

### RNA isolation and cDNA synthesis

Cell lysate was used for automated RNA isolation using a KingFisher Apex instrument with a 96 deep-well head, following the manufacturer’s protocol. Residual DNA was removed using the Lucigen DNase kit (Cat#D9905K) or TURBO DNase (Thermo Fisher, AM2238). Final RNA yields were measured using a Qubit instrument before proceeding to cDNA synthesis. cDNA was synthesized from 200 ng DNase-treated RNA using the ProtoScript II First Strand kit (NEB, #E6560L) with oligo-dT primers and random hexamers (1:1). RNA and primers were denatured at 95°C for 3 minutes, then reverse transcription was performed at 48°C for 1 hour followed by enzyme inactivation at 80°C for 5 minutes.

### Immunocytochemistry

Cells were fixed in 4% PFA/4% sucrose for 15 minutes at room temperature, then permeabilized and blocked in 0.2% Triton X-100 in 5% normal goat serum (NGS) for 30 minutes at room temperature. Primary antibodies were diluted in PBS containing 2% NGS and applied overnight at 4°C: anti-MECP2 (Cell Signaling Technology, #3456S; rabbit, 1:200) and anti-MAP2 (Thermo Fisher, #PA1-10005; chicken, 1:10,000). Plates were sealed with parafilm during incubation. Cells were then washed three times with PBS and incubated with secondary antibodies diluted 1:500 in PBS containing 2% NGS for 1 hour at room temperature. Secondary antibodies used were goat anti-rabbit IgG Alexa Fluor 647 (Thermo Fisher, #A-21244) and goat anti-chicken IgG Alexa Fluor 488 (Thermo Fisher, #A-11039). Nuclei were counterstained with DAPI (1:100 in PBS) for 10 minutes at room temperature. Cells were washed three times with PBS, and plates were sealed and stored at 4°C in the dark until imaging on the Zeiss Axios Observer 7.

### Western Blot

Protein samples were prepared in Bolt 4X LDS Sample Buffer (Thermo Fisher, #B0007) and Bolt 10X Reducing Agent (Thermo Fisher, #B0009) and denatured at 95°C for 5 minutes before loading. *In vitro* samples were loaded at 20 µg total protein; brain homogenates were normalized to 50 µg and denatured at 95°C for 5 minutes before loading. Proteins were resolved on 4–12% NuPAGE Bis-Tris Midi gels (Thermo Fisher, #WBT41212BOX) at 200V for 40 minutes using MOPS running buffer. Proteins were transferred to nitrocellulose membranes using the iBlot 2 system (P0, 7 minutes; Thermo Fisher).

Membranes were blocked in Pierce Fast Blocking Buffer (Thermo Fisher), then incubated overnight at 4°C in blocking buffer with the following antibodies: anti-MeCP2 (CST #3456S for cell lysates at 1:1,000 or Invitrogen #49-1029 for brain homogenate at 1:1,000) and anti-GAPDH (BioLegend, #649202 for cell lysates at 1:10,000 or Abcam, ab181602 for brain homogenate at 1:10,000). Membranes were washed three times in 1X TBS-T (5 minutes each) and incubated for 1 hour at room temperature with HRP-conjugated secondary antibodies in blocking buffer: goat anti-rabbit IgG (1:10,000; LI-COR, #926-80011) and, for *in vitro* samples only, mouse IgGκ BP-HRP (1:5,000; Santa Cruz Biotechnology, #sc-516102). After three additional TBS-T washes, *in vitro* membranes were incubated with ECL substrate for 5 minutes prior to imaging. All membranes were imaged on an iBright imaging system (Thermo Fisher).

### NGS Library Preparation

Three biological replicates were generated per condition across two independent differentiations (48 samples total; 6 Vehicle replicates per genotype). For each biological replicate, three wells per condition were pooled prior to RNA extraction. RNA integrity was confirmed by TapeStation (RIN ≥ 8.5). Sequencing libraries were prepared from 500 ng total RNA using the NEBNext Ultra II Directional RNA Library Prep Kit for Illumina with poly(A) mRNA selection (NEBNext Poly(A) mRNA Magnetic Isolation Module). Libraries were amplified for 15 PCR cycles using NEBNext 96 Unique Dual Index primers, purified by SPRI bead cleanup, and pooled at equimolar ratios. Paired-end sequencing (2 × 150 bp) was performed by Azenta on an Illumina NovaSeq platform targeting a minimum of 100 million reads per sample.

### RNA-seq Read Processing and Alignment

Sequencing reads were preprocessed with fastp (v0.22.0) to remove adapters (automatic paired-end detection), trim poly-G tails, and filter reads shorter than 50 bp. Trimmed reads were aligned to the human reference genome (GRCh38.p14, Gencode v47) using STAR (v2.7.9a) with a custom index augmented with AAV plasmid sequences to prevent mis-mapping of gRNA-derived reads to the endogenous *MECP2* locus. Alignment was performed with default RNA-seq parameters including splice junction detection, and transcriptome-aligned BAM output was generated for downstream quantification. PCR duplicates were marked using GATK MarkDuplicates (v4.3.0.0), and overall alignment quality was assessed with MultiQC (v1.14).

### Contrast Design

A unified contrast structure was applied across gene expression, global differential splicing, and differential RNA editing analyses, with all comparisons stratified by genotype (R168X or WT) and dose (5 × 10^4^ or 5 × 10^5^ vg/cell). For primary off-target assessment, each payload (MECP2-03 or MECP2-04) was compared against dose- and genotype-matched NTGC controls (n = 3 biological replicates per group). AAV backbone effects were characterized by comparing NTGC against Vehicle controls (n = 3 vs. 6 replicates per genotype); events significant in these contrasts were excluded from payload-specific off-target counts to isolate gRNA-dependent effects. Cross-genotype contrasts (e.g., MECP2-03 R168X vs. NTGC WT) were included to evaluate whether restoration of MECP2 expression shifts the R168X transcriptome toward a WT state.

### Gene Expression Analysis

Transcript-level quantification was performed with RSEM (v1.3.3) using transcriptome-aligned BAMs from STAR, and expected gene-level counts were used as input to PyDESeq2 (v0.4.4) for differential expression analysis. Genes with a Benjamini-Hochberg adjusted p-value < 0.05 and |log_2_ fold change| > 1.0 were considered differentially expressed. For AAV backbone contrasts (NTGC vs. Vehicle), a more permissive threshold of |log_2_FC| > 0.75 was applied to maximize exclusion of vector-induced expression changes from payload-specific off-target counts

### Global Differential Splicing Analysis

Transcriptome-wide alternative splicing was quantified with rMATS (v4.1.2 turbo) using coordinate-sorted BAMs, covering five event types: skipped exon (SE), alternative 5’ splice site (A5SS), alternative 3’ splice site (A3SS), mutually exclusive exons (MXE), and retained intron (RI). Splicing events were considered significant if they met thresholds of FDR < 0.05, |ΔPSI| ≥ 0.25, and junction read depth > 100. For near-constitutive exons (baseline PSI > 0.9 or < 0.1), a more sensitive threshold of |ΔPSI| ≥ 0.10 was applied, as even modest changes in inclusion at these highly constrained positions may reflect biologically meaningful perturbations. For AAV backbone contrasts, thresholds were relaxed (read depth > 20, |ΔPSI| ≥ 0.08) to maximize sensitivity for vector-induced splicing changes.

### Differential RNA Editing Analysis

Site-level differential A-to-I editing was assessed with JACUSA2 (v2.0.4), which compares base composition between treatment and control samples at each genomic position using a Dirichlet-Multinomial log-likelihood ratio test. Strand-specific analysis was performed to distinguish A-to-G changes on the plus strand from T-to-C changes on the minus strand (both indicative of A-to-I editing). Known SNPs from gnomAD and the 1000 Genomes Project were masked to prevent false-positive calls. Sites in intergenic regions were excluded to focus on functional loci. Candidate off-target editing sites were filtered to require a JACUSA2 score ≥ 10 (equivalent to p < 0.001 by chisquared approximation), a positive Δ(A-to-G) > 0.05 (consistent with ADAR-mediated editing), mean read depth > 50 across treatment replicates, and a control baseline A-to-G frequency < 0.05 to exclude fluctuations at endogenous editing sites. Sites significant in AAV background contrasts (NTGC vs. Vehicle) were excluded to remove AAV transduction artifacts.

### *In Silico* Off-Target Prediction

Potential gRNA hybridization sites were predicted by aligning the MECP2-03 and MECP2-04 gRNA sequences against the human genome (GRCh38.p14) using BLAST+ (v2.13.0) with sensitive parameters (word size 10, e-value 100,000). Hits were filtered to require ≥80% sequence identity over ≥25 bp and were annotated against Gencode v47 gene models using bedtools (v2.30.0) with strand-aware intersection to ensure predicted hits are antisense to the transcribed sequence. Hits within exons or within 500 bp of exon boundaries were retained. Because gRNAs expressed from an snRNA scaffold localize to the nucleus, analysis was restricted to the pre-mRNA transcriptome.

### Off-Target Classification

Off-target events identified by functional assays were classified as hybridization-dependent or hybridization-independent based on overlap with BLAST-predicted gRNA binding sites. Classification criteria were tailored to each assay: for editing events, the corresponding BLAST hit was required to directly overlap the edit site; for splicing events, the hit was required to fall within 500 bp of any exon involved in the event; and for gene expression changes, the hit was required to overlap any exon or exon-proximal region (within 500 bp) of the affected gene. Events passing significance thresholds at the high dose were retained; events significant only at the low dose were included only if the same event was independently significant at the high dose with a consistent direction of effect.

### Animals

*Mecp2*^R168X^ mice, stock #019528, were purchased from Jackson Laboratory and a live colony maintained by crossing heterozygous females (*Mecp2*^R168X/+)^ with wild-type *Mecp2*^+/y^ males. Mice were housed up to 5 per cage and kept on a 12-h light–dark cycle with ad libitum food (Teklad/Envigo, Global 19% Protein Extruded Rodent Diet, Irradiated, Cat# 2919) and water. Vivarium temperature was controlled between 68–79 °F with relative humidity maintained at 30– 70%. Mice were genotyped as previously described^16^ and *Mecp2*^R168X/y^ males were randomized into one of four treatment groups: vehicle or one of three AAV-delivered gRNA payloads (MECP2-02, MECP2-03, or MECP2-04) across two cohorts. A single retro-orbital injection was administered between postnatal days 14–19, and animals in the molecular cohort (n=5 per group) were euthanized by CO2 at four weeks post-treatment and brains were extracted and sagittally bisected for downstream processing. Animals in the survival cohort (n=10 per group) were monitored over a 184-day observation period for survival, body weight, and Rett phenotype scores. Animals remaining at the end of the 184-day observation period were sacrificed and brains were extracted for downstream processing.

Experimental procedures were approved by the Institutional Animal Care and Use Committee (IACUC) at Omeros Corp under protocol #24-01.

### Tissue Processing

Grinding vials containing the right hemisphere (excluding the cerebellum and olfactory bulbs) were placed in a 24-well aluminum cryo-block (SPEX SamplePrep cat# 2663) in a bath of liquid nitrogen prior to grinding for improved shattering. Samples were cryogenically ground on the Mini-G homogenizer (SPEX SamplePrep, cat# 1600) at a rate of 1500 rpm for 1 minute to obtain a homogenous powder. Homogenized powder was separated for RNA and protein isolation. For RNA isolation, cells were lysed in 500 µL MagMAX lysis/binding buffer containing 0.7% β-mercaptoethanol and processed with 100 µL input per well. Eluted RNA was treated with Lucigen DNase I (1.5× reaction: 25.5 µL RNA, 1.5 µL enzyme, 3 µL 10× buffer) for 30 minutes at room temperature, then inactivated at ™80°C. RNA concentration was measured by Qubit fluorometry (High Sensitivity RNA assay). Remaining powder was lysed in 1ml of 1X RIPA buffer containing HALT protease and phosphatase inhibitors.

### Immunohistochemistry

The left brain hemisphere was fixed in 10% neutral buffered formalin for 24-48 hours and paraffin embedded at the University of Washington histology core. 5 µm sections were stained for Mecp2 using automated immunohistochemistry on the Leica Bond RX automated immunostainer. Slides were deparaffinized using Bond Dewax Solution (Leica, #AR9222) and subjected to heat-induced epitope retrieval in Bond Epitope Retrieval Solution 1 (citrate buffer, pH 6.0; Leica, #AR9961) for 10 minutes at 100°C. Sections were blocked for 30 minutes at room temperature in blocking buffer consisting of 10% normal goat serum (Jackson ImmunoResearch, #005-000-121) and 1% BSA in PBS-T (1X PBS, 0.1% Tween-20). The primary antibody against Mecp2 (D4F3; Cell Signaling Technology, #3456; rabbit monoclonal) was diluted 1:400 in Bond Primary Antibody Diluent (Leica, #AR9352) and applied for 90 minutes at room temperature via two dispense steps for 30 min and 60 min respectively. Detection was performed using a goat anti-rabbit nanobody conjugated to Alexa Fluor 647 (Fluo-tag, N2404-AF647-S, 1:500 in 1% BSA/PBS-T) applied for 60 minutes at room temperature via two dispense steps, each for 30 minutes. Nuclei were counterstained with DAPI for 5 minutes at room temperature and slides were counterstained with hematoxylin (Leica, DS9800). Slides were mounted with aqueous mounting medium. Stained sections were imaged on the Zeiss Axios Observer 7 and quantified using Zeiss Zen Blue Image Analysis Software.

### Data Analysis

Percent RNA editing at the *MeCP2*^R168X^ target site, bulk protein restoration, % MeCP2 positive, and median fluorescence intensity (MFI) was quantified from representative FOVs from cortex and midbrain. Pairwise comparisons of each treatment versus vehicle were performed using two-sided Mann-Whitney U tests, with full pairwise comparisons among all four groups (six pairs) subjected to Holm correction.

### Survival Analysis

Animals euthanized for intercurrent, non-Rett-related causes (n = 4; tail wounds, malocclusion) were right-censored at the date of euthanasia. Animals alive at the pre-specified study termination date (September 19, 2025; n = 5) were right-censored at that date. All data collected prior to censoring were included in the analysis. Survival was defined as time from injection to death or censoring. Kaplan-Meier survival curves were estimated for each treatment group with at-risk tables reported at 30-day intervals, and median survival times with 95% confidence intervals were extracted from the Kaplan-Meier estimator. Pairwise comparisons of each treatment versus vehicle were conducted using the log-rank test, with p-values corrected for three simultaneous comparisons using the Holm method. Full pairwise comparisons among all four groups (six pairs) were performed using the log-rank test, penalized Cox proportional hazards regression, and restricted mean survival time (RMST), with Holm correction applied within each test. Hazard ratios were estimated by pairwise Cox proportional hazards regression. Given the small per-group sample size (n = 9–10) and near-complete separation in the MECP2-04 arm (4 of 10 events observed), a penalized Cox model (L2 penalty, λ = 0.1) was used as the primary analysis to ensure stable coefficient estimates; unpenalized Cox regression was performed as a sensitivity analysis. The proportional hazards assumption was assessed using Schoenfeld residuals. As an assumption-free complement to Cox regression, RMST was computed at τ = 184 days for each group, with differences versus vehicle estimated using 95% bootstrap confidence intervals (10,000 resamples).

### Longitudinal Phenotype Analysis

Rett phenotype scores, including a composite score and five sub-components (mobility, gait, hindlimb clasping, tremor, and general condition), and body weight were assessed longitudinally. To mitigate survivor bias arising from earlier mortality in vehicle-treated animals, longitudinal analyses were restricted to a data-driven window (days 0–43 post-treatment), defined as the last assessment day on which at least five vehicle-treated animals contributed phenotype data. The primary longitudinal model was a linear mixed-effects (LME) model of the form: **Score ~ Tracking_day × Treatment + (1** | **Animal)** where Treatment was dummy-coded with vehicle as the reference group and animal was included as a random intercept to account for repeated measures. The treatment-by-time interaction term was the primary coefficient of interest, estimating the difference in daily rate of phenotype change between each treatment and vehicle; negative values indicate slower phenotypic progression relative to vehicle. As a sensitivity analysis, generalized estimating equations (GEE) with an exchangeable correlation structure were fitted to the same data. Model fit was evaluated against a GEE model with first-order autoregressive (AR(1)) correlation using the quasi-likelihood under the independence model criterion (QIC), and the better-fitting model was reported. Full pairwise inter-treatment comparisons of longitudinal trajectories were conducted by refitting the LME model for each group pair, with Holm correction applied across all six pairwise comparisons within each outcome.

### Software and Reproducibility

All *in vitro* analyses were executed within a custom Nextflow (DSL2) pipeline using versioned Docker containers. *In vivo* statistical analyses were performed in Python 3.12 using lifelines (v0.29) for survival analysis, statsmodels (v0.14) for LME and GEE models, and scipy (v1.14) for non-parametric tests. Figures were generated with matplotlib (v3.9) and seaborn (v0.13). Key software versions: fastp 0.22.0, STAR 2.7.9a, RSEM 1.3.3, GATK 4.3.0.0, rMATS 4.1.2, JACUSA2 2.0.4, BLAST+ 2.13.0, bedtools 2.30.0, samtools 1.14, MultiQC 1.14, PyDESeq2 0.4.4.

## Acknowledgements

The authors gratefully acknowledge Joanne Boysen, Lauren Swanson, and Aditya Radhakrishnan for their contributions to in-cell editing pre-processing analysis, Lynsey Kovar for their support with specificity assessment and Bora Banjanin, Aydin Abiar, and Ron Hause for their work on generative gRNA design. We thank Alex Rohde, Ben Weigler, and Miguel Barrera (Omeros Corp) for their assistance with phenotype scoring and animal monitoring. We are grateful to the Rett Syndrome Research Trust (RSRT) and System 1 Biosciences for providing iPSCs, and to Katherine Stewart and Turnee Malik for their contributions to rational gRNA screening. We thank Brooke Cichelli, Leslie Blakely, and Jonathan Wang for colony management, sample processing, animal monitoring, and support with Rett Phenotype Scoring. We also thank Erin Gaffney for intellectual property support, Jennifer Moon for iPSC neuron support, Chelsea Fortin for cell imaging support, and Crystal Pontrello for primary neuron culture support. Virus production was supported by Haili Adams, Rebecca Olson, Indraneel Salukhe, Dhyani Patel, Kara Graves, Kimberly Franck, and Zaineb Alkadban. Virus purification was carried out by William Johnsen and Ashley Benson, and virus characterization was performed by Jeff Dantzler and Emma Thuline. The authors extend their sincere appreciation to all contributors for their essential roles in this work.

## Author contributions

Yiannis A. Savva, Brian J. Booth, Melissa G. Works, David J. Huss, Adrian W. Briggs, and Alison A. VanSchoiack conceived the study. Lucia Shumaker designed, led, and executed in vitro and in vivo experiments under the supervision of Alison A. VanSchoiack. Rachael Fasnacht led and executed in vivo experiments under the supervision of Eric M. Chadwick. Stephen M. Burleigh led vector optimization efforts under the supervision of Collin Hauskins and Yiannis A. Savva. Yue Jiang, Yingxin Cao, Lina R. Bagepalli, Andrew Sadowski, and Scott Rich designed and validated AI-generated gRNAs under the supervision of Brian J. Booth and Yiannis A. Savva. Brian Johnson conducted immunohistochemistry experiments under the supervision of Alison A. W. VanSchoiack. Forrest Golic and Nicole Enger performed screens for vector optimization under the supervision of Stephen M. Burleigh. Rachel Feiring generated experimental reagents and established protocols. Anupama Lakshmanan led reporter construct development and cell line engineering. Nikita Milani sup-ported in vitro and in vivo experiments. Yiannis A. Savva, Brian J. Booth, Yue Jiang, and Alison A. VanSchoiack wrote the manuscript with input from all authors. All authors reviewed and approved the final manuscript.

## Competing interests

All authors are current or former employees of Shape Therapeutics, Inc. who are inventors on patents and/or patent applications based on the site-directed RNA editing methods and/or the machine learning models described in this work. The authors declare no other competing interests.

