## Supplementary Figures for "Rett syndrome lifespan extension in mice via AI-guided ADAR editing"

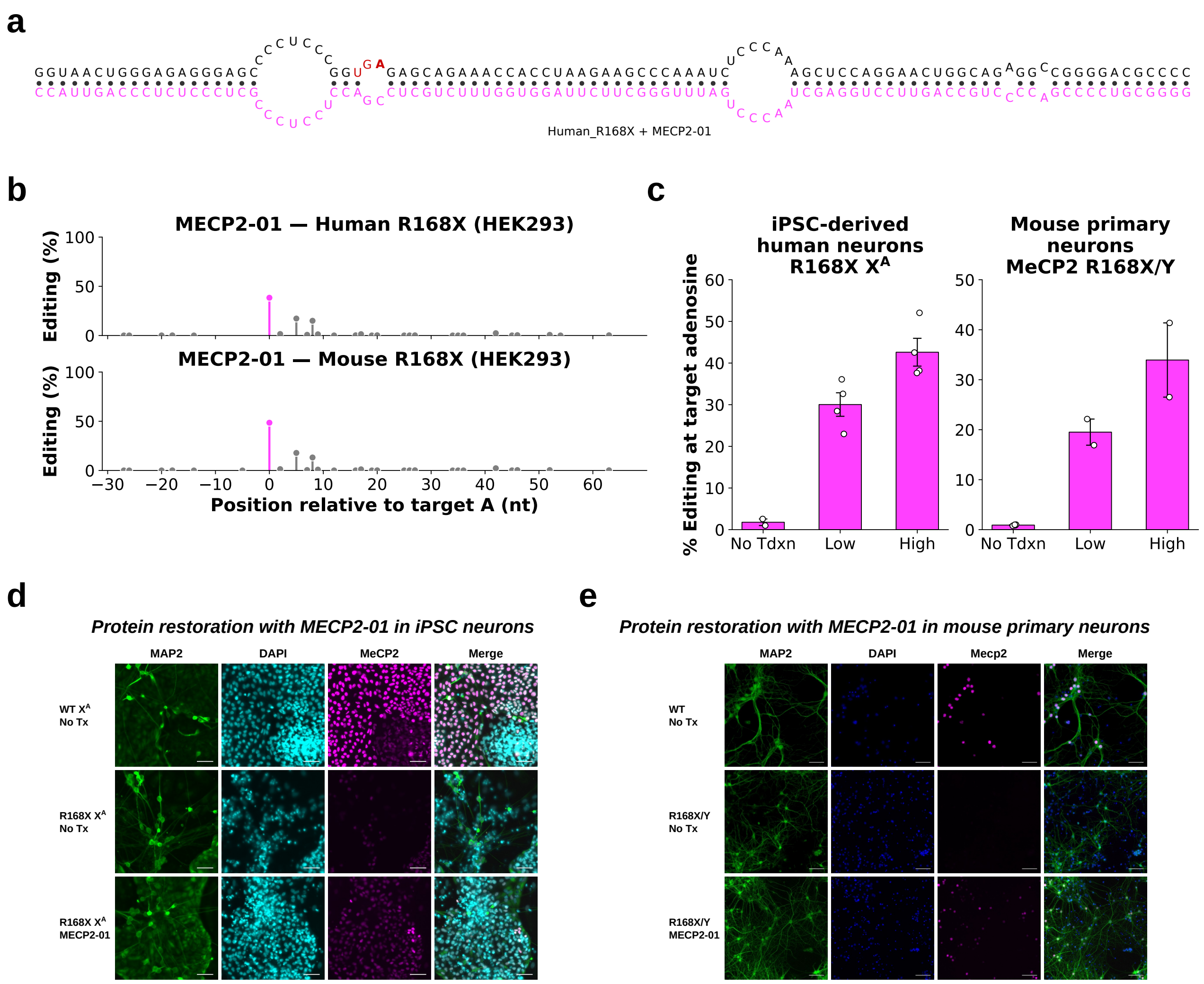


**Supplementary Fig. 1: A rational gRNA enables RNA editing and MeCP2 protein restoration in human and mouse neuronal models.**

**a**, Secondary RNA structure of an ADAR-recruiting rationally designed gRNA, MECP2-01, hybridized to the target *MECP2* R168X transcript. The gRNA (magenta) forms a duplex with the target RNA, positioning the adenosine-to-inosine (A-to-I) editing site (highlighted in red) in a favorable structural conformation. Bulges and mismatches are incorporated into the gRNA design to enhance ADAR recruitment and enhance editing selectivity. **b**, Local A-to-I editing profile of MECP2-01 in HEK293 cells expressing either the human R168X (top) or mouse R168X (bottom) *MECP2* transcript. Editing efficiency (%) is plotted against nucleotide position relative to the target adenosine (position 0, magenta). **c**, Quantification of on-target A-to-I editing efficiency at the R168X adenosine in patient iPSC-derived neurons carrying the R168X mutation (left) and mouse primary neurons from MeCP2 R168X/Y mice (right), following transduction with low or high doses of MECP2-01 gRNA and no transduction (No Tdxn) control. Bar graphs represent mean $\pm$ SEM; individual data points are shown. Immunofluorescence analysis of MeCP2 protein restoration in iPSC-derived human neurons (**d**) and mouse primary neurons (**e**) from wild-type (WT), R168X mutant, and R168X mutant treated with MECP2-01 gRNA. Cells were stained for the neuronal marker MAP2 (green), nuclear marker DAPI (blue), and MeCP2 (magenta). Composite images are shown in the rightmost column. Scale bars = 50µm.

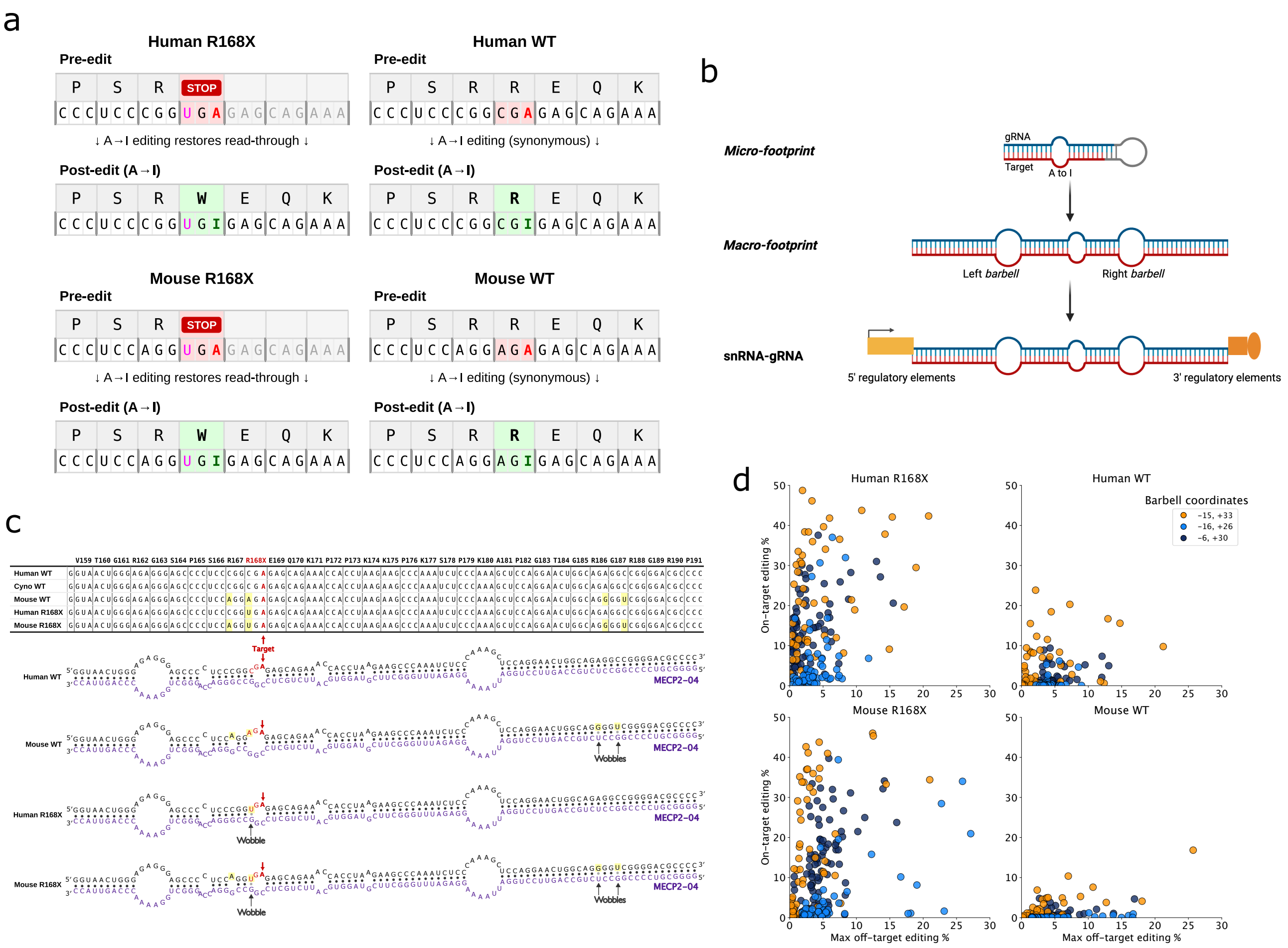


**Supplementary Fig. 2: Design and optimization of ADAR-recruiting gRNAs across human and mouse sequence contexts.**

**a**, Schematic of target codons and predicted editing outcomes for human and mouse *MECP2* sequences in R168X and wild-type (WT) contexts. Pre-edited sequences (top) depict the R168X allele harboring a UGA premature termination codon (red) and the WT allele harboring a CGA arginine codon. Adenosine-to-inosine (A-to-I) editing at the target adenosine (highlighted in green; post-edit) converts UGA to UGI (inosine read as guanosine, interpreted as tryptophan (W) thereby restoring translational read-through in the R168X context. In the WT context, editing at the equivalent position results in a synonymous CGA-to-CGI codon change (arginine, R). Predicted amino acid identities are indicated above each codon. **b**, Schematic illustrating the hierarchical gRNA engineering strategy. An initial micro-footprint, a minimal antisense hybridization domain, is first optimized in a high-throughput biochemical assay. The optimized micro-footprint is subsequently incorporated into a macro-footprint design comprising an extended hybridization region flanked by 5′ and 3′ barbell structures, which promote ADAR recruitment and enhance editing efficiency. The macro-footprint is then embedded within a snRNA expression framework, flanked by cognate 5′ and 3′ regulatory elements to confer nuclear localization, gRNA stability, and robust expression. **c**, (Top) Multiple sequence alignment of human WT, cynomolgus macaque (Cyno) WT, mouse WT, human R168X, and mouse R168X *MECP2* target region sequences. The target adenosine is indicated by a red arrow; nucleotide positions diverging from the human WT reference are highlighted in yellow. (Bottom) Predicted RNA secondary structures of the MECP2-04 gRNA for each sequence context (Human WT, Mouse WT, Human R168X, Mouse R168X), with the target adenosine (red arrow), wobble mismatch positions, and the gRNA sequence designation (purple) annotated for each structure. **d**, Scatter plots depicting on-target editing efficiency (%; y-axis) versus maximum off-target editing frequency (%; x-axis) for a panel of micro-footprint variants embedded within three barbell coordinate configurations and evaluated across four sequence contexts: Human R168X, Human WT, Mouse R168X, and Mouse WT. Each data point represents an individual macro-footprint construct, colored according to barbell coordinate position: (−15, +33), (−16, +26), or (−6, +30).

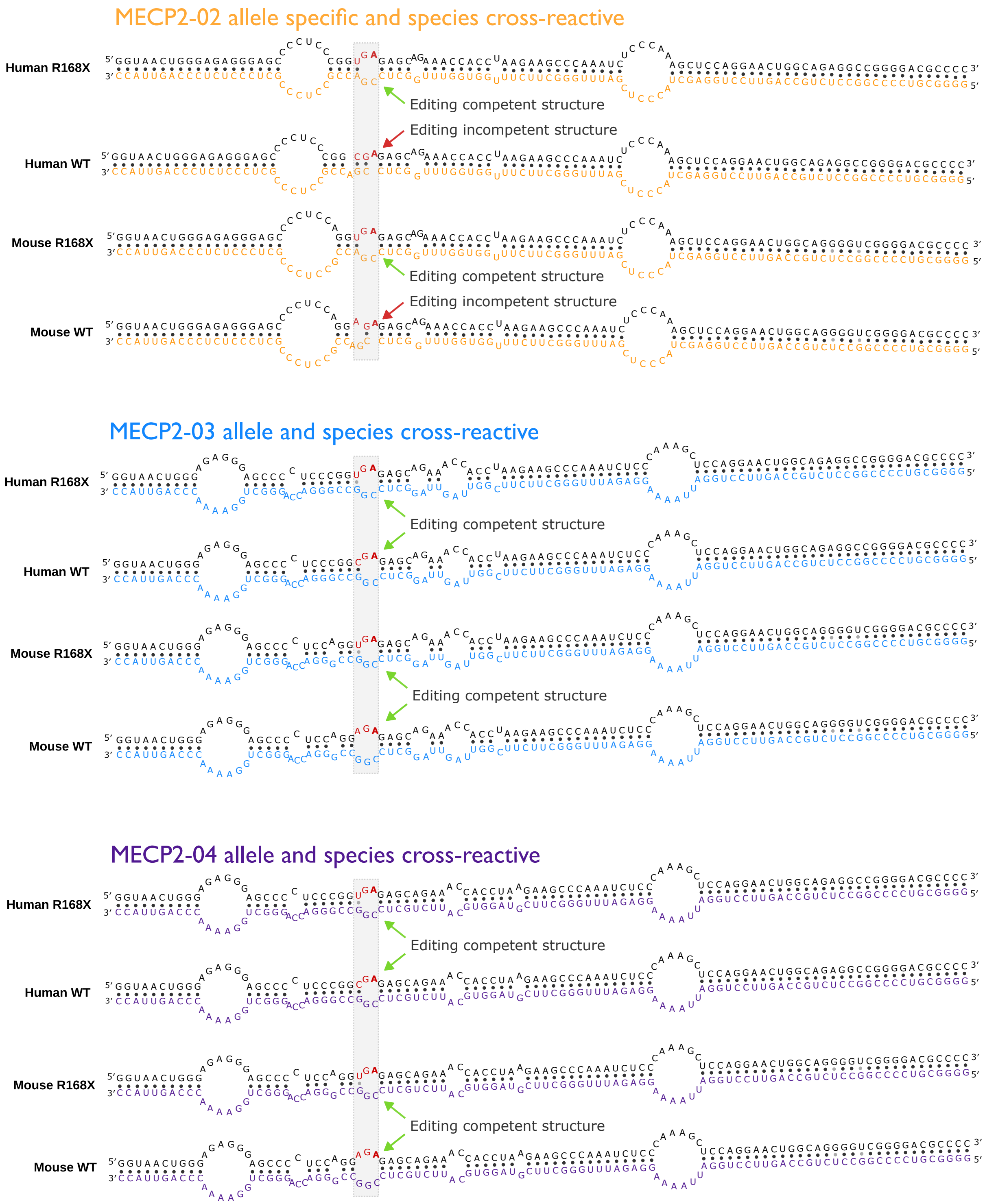


**Supplementary Fig. 3: RNA secondary structure predictions for top *MECP2*-targeting gRNAs across human and mouse alleles.**

Predicted RNA secondary structures in human and mouse *MECP2* transcripts (grey) for the top three gRNA designs. Each panel depicts folded RNA structures for the R168X mutant and WT sequences in both human and mouse. The target adenosine is indicated in red. **MECP2-02** (Top, orange), an allele-specific and species cross-reactive gRNA design. R168X mutant transcripts adopt an editing-competent structure (green arrow), while WT transcripts adopt an editing-incompetent structure (red arrow), enabling preferential editing of the disease allele. This allele-selective behavior is conserved across human and mouse sequences. **MECP2-03** (Middle, blue) and **MECP2-04** (Bottom, purple)**,** are non-allele specific gRNAs that are species cross-reactive. Both human and mouse R168X mutant transcripts adopt an editing-competent structure (green arrows). WT transcripts are predicted to adopt slightly alternative structures that also support efficient editing. Editing-competent structures are defined by accessibility of the target adenosine within the local folding context required for ADAR-mediated deamination. Structures were predicted using Vienna fold RNA duplex with default parameters.

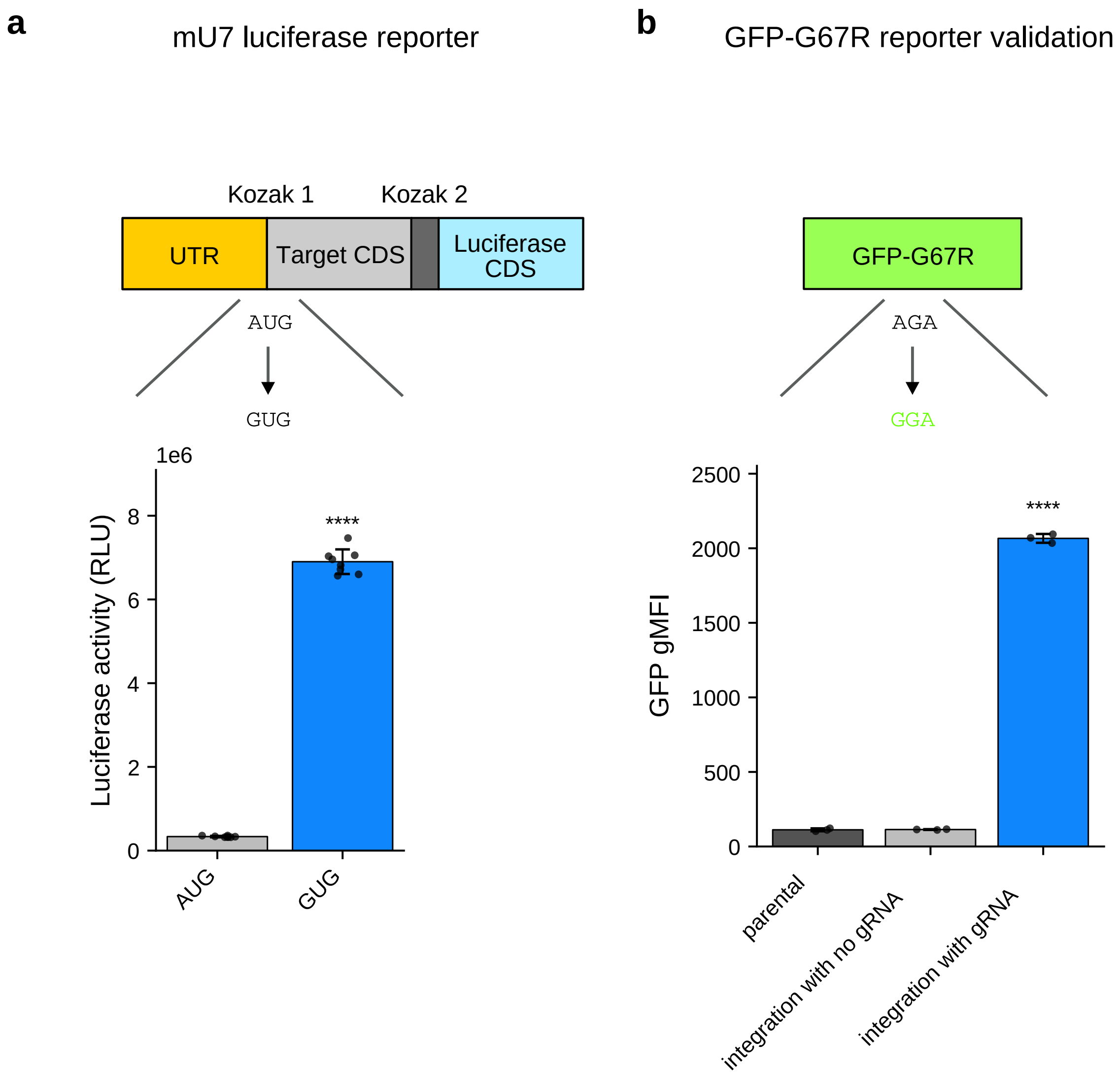


**Supplementary Fig. 4: Reporter constructs for monitoring adenosine-to-inosine deamination activity.**

**a,** A start codon-based reporter system was developed to detect deamination of AUG codons. A secondary luciferase reading frame was inserted downstream and out-of-frame relative to a test codon, such that translation initiating from the upstream codon suppresses luciferase expression. Deamination of the start codon (AUG→IUG, read as GUG) disrupts canonical translation initiation, enabling ribosomal access to the downstream luciferase frame. Validation is demonstrated by comparison of constructs harboring either an AUG or GUG start codon: the GUG construct yields high luciferase activity, consistent with impaired upstream initiation, whereas the AUG construct produces minimal signal. This system is hereafter referred to as the "Kozak competition" reporter. **b,** A GFP-based reporter was generated to detect deamination of a specific adenosine within the coding sequence. Introduction of a G67R missense mutation (GGA→AGA) renders GFP non-fluorescent. Adenosine-to-inosine deamination at this position converts the AGA codon back to an inosine-containing codon read as GGA, thereby restoring the glycine residue at position 67 and rescuing GFP fluorescence. This reporter thus provides a direct readout of site-specific A-to-I editing activity. Bars represent mean ± SEM; two-sided Welch's t-test vs control: *$p<0.05$, **$p<0.01$, ***$p<0.001$, ****$p<0.0001$.

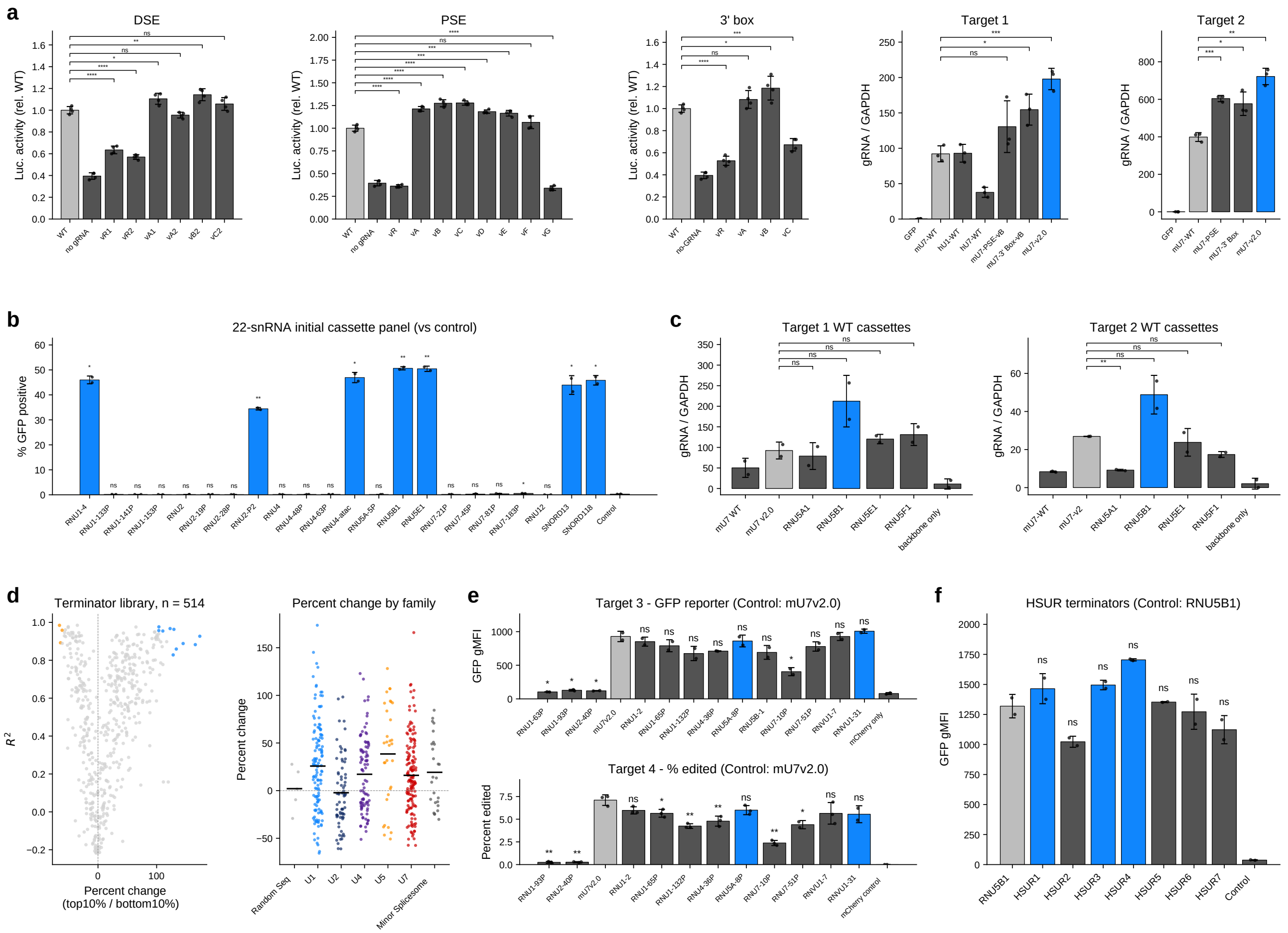


**Supplementary Fig. 5: Optimization of snRNA regulatory elements and terminator sequences for enhanced guide RNA expression.**

**a,** Systematic mutagenesis of snRNA cis-regulatory elements affecting guide RNA expression. Luciferase reporter assays, using the kozak competition reporter, measuring the effect of mutations in the distal sequence element (DSE, left), proximal sequence element (PSE, middle), and 3' box (right) on transcriptional activity, shown relative to wild-type (WT). ddPCR quantification of guide RNA levels normalized to GAPDH for two gRNAs for non-MeCP2 validation targets, Target 1 (fourth panel) and Target 2 (fifth panel), across constructs harboring key regulatory element variants. **b,** Flow cytometry-based screen of an initial 22-snRNA cassette panel measuring percent GFP-positive cells. Blue bars indicate cassettes with statistically significant activity relative to the control condition. **c**, ddPCR quantification of guide RNA expression (normalized to GAPDH) from wild-type snRNA cassettes for Target 1 (left) and Target 2 (right) across selected RNU5 cassettes. **d**, Terminator library screen (n = 514). Scatter plot (left) showing the correlation coefficient ($R^2$) versus percent change (top 10%/bottom 10%) for all terminator variants. High-performing candidates are highlighted in light blue color, while low performing sequences are highlighted in orange color. Dot plot (right) showing percent change in guide RNA activity grouped by snRNA family (Random Seq, U1–U7, Minor Spliceosome), with colored dots representing individual terminator families. **e**, Functional validation of top terminator candidates using a GFP reporter for Target 3 (top, GFP gMFI, control: mU7v2.0) and percent editing for Target 4 (bottom, control: mU7v2.0). Blue bars highlight the two lead candidates while the light grey bars reference mU7v2.0 control construct. **f**, Evaluation of HSUR (Herpesvirus saimiri U-rich) terminators using GFP gMFI as readout (control: RNU5B1). Individual HSUR variants (HSUR1–HSUR7) are compared to the RNU5B1 control (light gray) and a negative control. For all panels, error bars represent SEM; Statistical comparisons: two-sided Welch's t-test vs reference column; *p<0.05, **p<0.01, ***p<0.001, ****p<0.0001, ns = not significant.

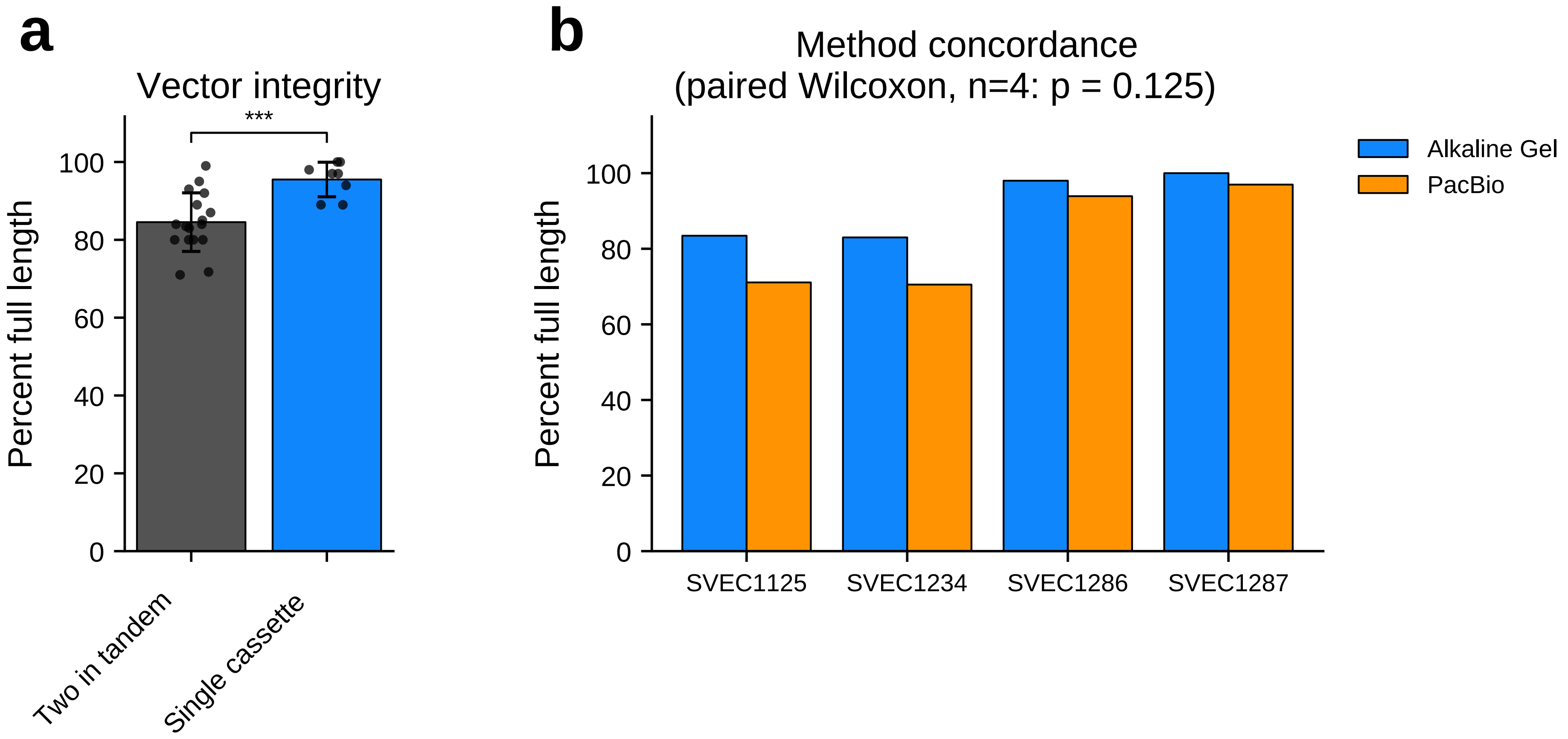


**Supplementary Fig. 6: Vector integrity and method concordance for single cassette and tandem AAV constructs.**

**a, Vector integrity.** Percent full-length vector genomes for "Two in tandem" (gray) and "Single cassette" (blue) AAV constructs, as measured by alkaline gel analysis. Individual data points are overlaid on each bar. *** $p < 0.001$. **b, Method concordance.** Comparison of percent full-length vector genome measurements obtained by alkaline gel electrophoresis (blue) versus PacBio long-read sequencing (orange) across four AAV vector lots (SVEC1125, SVEC1234, SVEC1286, and SVEC1287). No statistically significant difference was observed between the two quantification methods (paired Wilcoxon signed-rank test, $n = 4$; $p = 0.125$), supporting the concordance of alkaline gel analysis with orthogonal sequencing-based assessment of vector integrity.

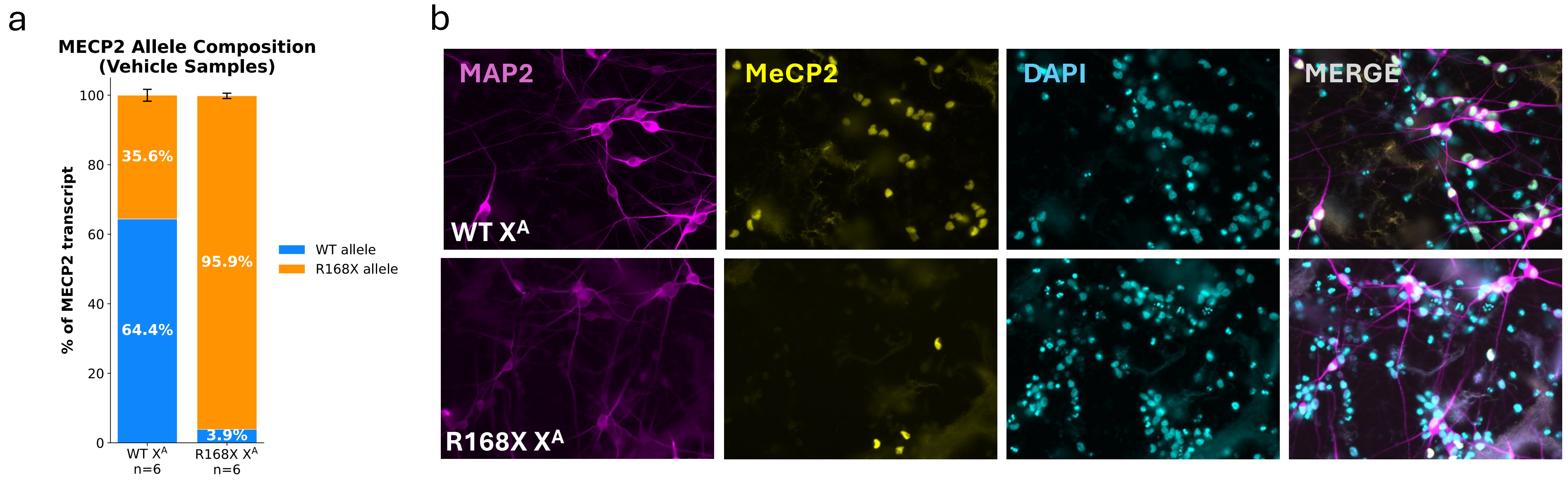


**Supplementary Fig. 7:** ***MECP2*** **allele expression and MeCP2 protein levels in WT and R168X iPSC-derived neurons.**

**a,** Stacked bar graph showing the relative proportions of *MECP2* transcript alleles in neurons from WT $X^A$ (n=6) and R168X $X^A$ (n=6) samples. The WT allele is shown in blue and the R168X allele in orange. Error bars represent SEM. **b,** Representative immunofluorescence images of WT $X^A$ (top row) and R168X $X^A$ (bottom row) neurons stained for the dendritic marker MAP2 (magenta), MeCP2 protein (yellow), and the nuclear marker DAPI (cyan), with a merged composite shown in the rightmost panel. WT $X^A$ neurons display robust MeCP2 immunoreactivity in nuclei, consistent with normal protein expression. R168X $X^A$ neurons show markedly reduced MeCP2 signal, reflecting loss of functional protein due to the nonsense mutation. MAP2 staining confirms the presence of neurons across all conditions, and DAPI labels all cell nuclei.

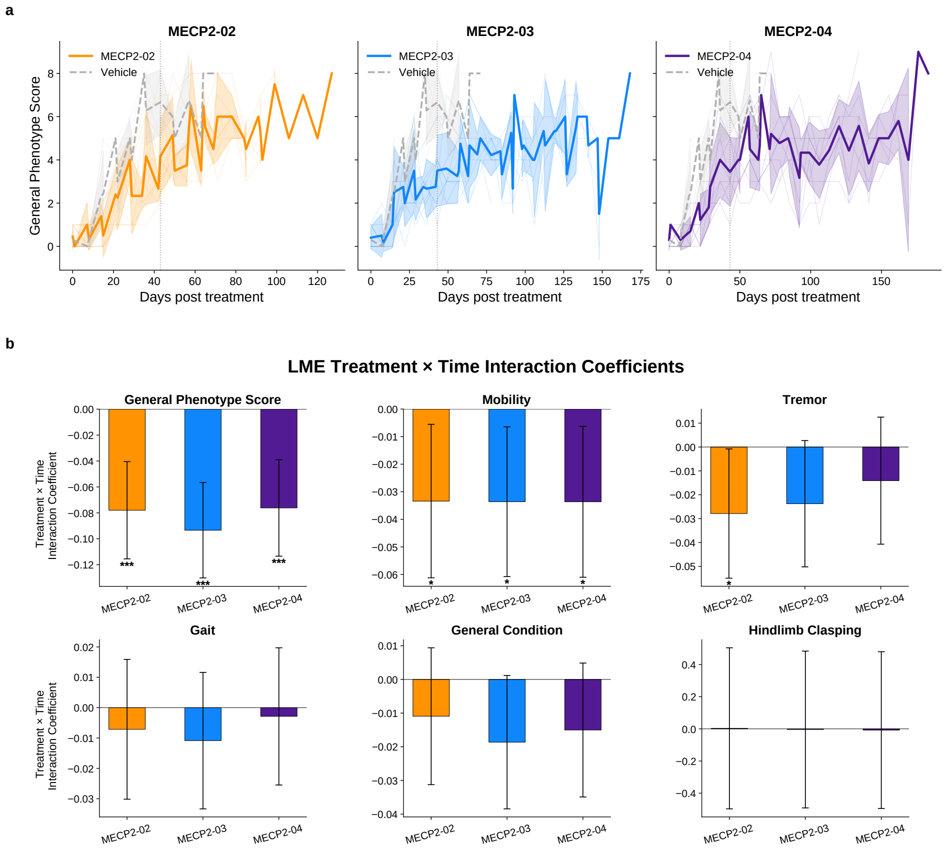


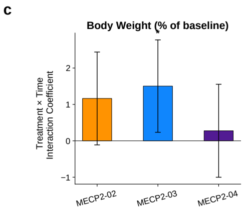


d

**Survival Analysis Summary — MECP2$^{R168X}$ Mouse Model**

Pairwise comparisons vs Vehicle (penalized Cox PH, penalizer=0.1)

| | Enrollment | | | Survival Endpoints | | | |
|---|---|---|---|---|---|---|---|
| Treatment | N | Events | Censored | Median Survival (days) | HR (95% CI) | RMST Difference (95% CI) | Log-rank p |
| Vehicle | 10 | 9 | 1 | 50 | 1.00 (ref) | — (ref) | — |
| MECP2-02 | 9 | 9 | 0 | 97 | 0.12 (0.03–0.49) | +39 (23–55) | 2.3e-04 |
| MECP2-03 | 10 | 8 | 2 | 126 | 0.09 (0.02–0.37) | +74 (50–98) | 5.7e-05 |
| MECP2-04 | 10 | 4 | 6 | 174 | 0.07 (0.02–0.35) | +115 (99–131) | 5.7e-05 |

HR = hazard ratio (penalized Cox PH). RMST = restricted mean survival time at τ = 184 days. Log-rank p-values are Holm-corrected for 3 comparisons. Censored animals include non-Rett euthanasia (n=4) and study-end survivors (n=5).

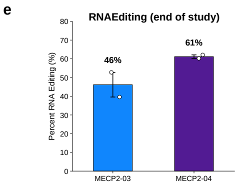


**Supplementary Fig. 8: Phenotypic, survival, and RNA-editing outcomes in the Mecp2$^{R168X}$ mouse model following gRNA treatment.**

**a,** Longitudinal general phenotype scores for each treatment cohort (MECP2-02, MECP2-03, MECP2-04) compared with vehicle controls (dashed, vehicle) over days post-treatment. Shaded bands represent ±1 SD. **b,** Linear mixed-effects (LME) model treatment × time interaction coefficients for six phenotypic domains: general phenotype score, mobility, tremor, gait, general condition, and hindlimb clasping. Bars represent coefficient estimates; error bars indicate 95% confidence intervals. Negative coefficients indicate slowed disease progression relative to vehicle. **c,** LME treatment × time interaction coefficient for body weight (% of baseline), showing change relative to vehicle over time. For b and c, asterisks denote significance (*$p<0.05$, **$p<0.01$,***$p < 0.001$). **d,** Survival analysis summary table for the Mecp2$^{R168X}$ mouse model. Pairwise comparisons versus vehicle were performed using penalized Cox proportional hazards regression (penalizer = 0.1). Columns report: treatment group, enrollment N, event and censored counts, median survival (days), hazard ratio (HR, 95% CI), restricted mean survival time (RMST) difference at $t = 184$ days (95% CI), and log-rank $p$-value (Holm-corrected for 3 comparisons). Censored animals include those euthanized for non-Rett causes ($n = 4$) and study-end survivors ($n = 5$). **e,** Percent RNA editing (A-to-I) at end of study for MECP2-03 and MECP2-04 gRNA treated cohorts, measured in target tissue. Bars show group means; error bars represent SEM.
