## Supplementary material for "Rett syndrome lifespan extension in mice via AI-guided ADAR editing": Table S1

| Off-Target Event Counts<br>MECP2-03 / MECP2-04 vs NTGC control | | | | | |
|---|---|---|---|---|---|
| Payload | Dose | vg/cell | Differential Expression | Differential Splicing | Differential A-to-G frequency |
| WT $X^A$ neurons | | | | | |
| MECP2-03 | low | 5E4 | 0 (0) | 0 (0) | 0 (0) |
| MECP2-04 | low | 5E4 | 0 (0) | 0 (0) | 0 (2) |
| MECP2-03 | high | 5E5 | 0 (0) | 0 (81) | 0 (283) |
| MECP2-04 | high | 5E5 | 0 (6) | 0 (46) | 0 (328) |
| R168X $X^A$ neurons | | | | | |
| MECP2-03 | low | 5E4 | 0 (0) | 0 (3) | 0 (0) |
| MECP2-04 | low | 5E4 | 0 (0) | 0 (0) | 0 (0) |
| MECP2-03 | high | 5E5 | 0 (0) | 0 (60) | 0 (322) |
| MECP2-04 | high | 5E5 | 0 (7) | 0 (17) | 0 (168) |

**Format:** X (Y) = hybridization-dependent (hybridization-independent) events

**DE thresholds:** padj < 0.05, |log2FC| ≥ 1.0 | **Splicing:** FDR < 0.05, |ΔPSI| ≥ 0.25 (or ≥ 0.10 at extreme baseline), depth ≥ 100 | **A-to-G:** Δ > 0.05, score ≥ 10

**Supplementary Table 1: Off-target events passing statistical and biological filters by RNAseq analysis.**

Table of off-target events that surpassed statistically and biological filters for differential expression, differential splicing, and differential A-to-G frequency by RNAseq (see **Methods**). The first number indicates hybridization dependent events and the number in parenthesis indicates hybridization independent events for each gRNA payload and genotype. Each MECP2-03 and MECP2-04 treated sample was contrasted with its dose matched NTGC negative control.
